# The neural correlates of novel versus familiar metaphors in healthy young adults: A functional near-infrared spectroscopy study

**DOI:** 10.1101/2025.10.12.681881

**Authors:** Anna Schwartz, Natalie M. Gilmore, Erin L. Meier

## Abstract

Despite extensive investigation, the neural correlates of metaphor processing remain debated. Poor theoretical and experimental control of variables that drive metaphor activation— particularly the constructs of novelty and familiarity—may be the reason for past discrepancies between studies. To address this issue, we used functional near-infrared spectroscopy (fNIRS) and a carefully designed paradigm modified from Cardillo et al. (2012) to investigate how neural activation varies by sentence type (metaphorical versus literal sentences) and novelty (completely novel versus familiarized phrases). Activity was significantly greater for metaphorical over literal sentences in the left inferior frontal gyrus, pars triangularis (LIFGtri), left inferior parietal cortex, right IFG, pars opercularis (RIFGop), and right angular gyrus (RAG). Novel metaphors to which participants had no prior exposure had significantly higher (albeit weak) effects within RIFGop, RAG, and right middle temporal gyrus (RMTG) compared to novel metaphors to which participants were exposed just prior to the fNIRS experiment. Pre-exposed, more familiar metaphors significantly activated a wider network of regions compared to novel metaphors, including bilateral middle frontal gyrus (MFG), bilateral IFGtri, and LMTG. A greater response time difference between conditions was associated with less LMFG activity for metaphors over literal sentences but higher LMTG activity for novel over more familiar metaphors. Taken together, these findings suggest that metaphors—particularly novel metaphors—do engage right hemisphere cortex more than other phrase types (literal sentences, more familiar metaphors) but that the effects are weaker than condition differences within canonical left language network and domain-general multiple demand network regions.

## 1. Introduction

Metaphor has been proposed as a mechanism that allows people to communicate novel (especially abstract) ideas to an audience, making them critical for learning and adapting to new situations throughout the lifespan (Di Paola et al., 2020; Gentner, 1983; Holyoak & Stamenković, 2018; Jamrozik et al., 2016; Lakoff & Johnson, 1980; Thibodeau & Boroditsky, 2011). Metaphors are considered the structural “backbone” of extensive cultural norms and heavily influential in structuring patterns of thought (Kövecses, 2010). Perhaps because of the ubiquity of metaphor use in adults’ everyday language, their neural correlates have been studied extensively, yet debate remains regarding which brain regions are specially recruited for metaphors versus other types of language, especially as metaphors are learned and become conventionalized.

### 1.1. Theories and evidence regarding the neural correlates of metaphor

This is particularly the case for conceptual metaphors, phrases in which two constructs are compared in order to highlight the aspect of one construct’s meaning (the target) that overlaps with another construct (the source). For example, the phrase THE JUDGE IS A JUGGLER uses juggling to highlight the way a judge (target) may need to simultaneously manage multiple tasks at the same time. The audience is expected to suppress the literal meaning of juggler (source), as a person who manipulates multiple physical objects, and instead apply the concepts of “simultaneous,” “manipulates,” and “multiple,” to the tasks of a judge. The conceptualization of the abstract link between the target and source domains is achieved by accessing pre-existing semantic and experiential knowledge of the source concept and mapping it to the target (Kövecses, 2010; Lakoff & Johnson, 1980). Because of this, theories regarding the neural underpinnings of metaphor have traditionally focused on hemispheric lateralization differences in semantic mapping involved in processing figurative language—including metaphors—versus literal phrases. For example, according to Jung-Beeman’s coarse semantic coding theory (Beeman, 1998; Beeman et al., 1994; Jung-Beeman, 2005), left hemisphere (LH) regions specialize in processing central word meanings and mapping closely-related concepts (i.e., fine coding)—as is necessary during literal sentence processing. On the other hand, within this theory, the right hemisphere (RH) has the capacity for fine coding in addition to processing peripheral or alternative word meanings and mapping concepts with distant relations (i.e., coarse coding), as is needed during metaphoric processing. Thus, according to the coarse semantic coding theory, literal language stimuli should recruit LH semantic network regions to a greater extent than their RH homologues whereas the opposite should be true of figurative language.

Systematic reviews and meta-analyses of prior neuroimaging investigations lend some credence to this LH versus RH distinction, but with mixed findings. Older meta-analytic studies (Bohrn et al., 2012; Rapp et al., 2012; Reyes-Aguilar et al., 2018; Yang, 2014) that included around 15 to 16 studies in their comparison of metaphoric and literal stimuli reported greater activation in the different parts of the right inferior frontal gyrus (IFG) (Brodmann areas [BAs] 44, 45, and 47) and right anterior to posterior superior temporal (STG) and middle temporal (MTG) gyri (BA 21 and 22) for metaphors over literal stimuli whereas minimal to no activation was seen in the RH for the opposite contrast (c.f. Bohrn et al., 2012). However, in a more recent meta-analysis of 30 studies, Huang et al. (2023) found that metaphors did not significantly activate any RH region to a greater extent than literal stimuli. Notably, all these meta-analyses also indicated that compared to literal stimuli, metaphors more greatly activated a wide network of left frontotemporal regions as well as several left medial and subcortical structures (e.g., medial prefrontal cortex [mPFC], anterior cingulate cortex [ACC]), but the specific areas varied by study. For instance, Bohrn et al. (2012), Yang (2014), and Huang et al. (2023) reported greater activity for metaphorical over literal stimuli in left IFG, pars orbitalis (IFGorb) and pars triangularis (IFGtri), whereas Rapp et al. (2012) found activation for this comparison only in LIFGorb, and Reyes-Aguilar et al. (2018) reported more posterior activation, specifically in LIFGtri and left IFG, pars opercularis (IFGop). As another example of mixed findings, the older reviews (Bohrn et al., 2012; Rapp et al., 2012; Yang, 2014) reported greater activation for metaphors in both anterior and posterior temporal regions whereas the more recent meta-analytic findings indicated greater metaphorical activity in posterior temporal regions only, specifically in posterior STG (pSTG) and MTG (pMTG) in Reyes-Aguilar et al. (2018) and pMTG extending into posterior inferior temporal gyrus (pITG) in Huang et al. (2023).

### 1.2. The effects of familiarity and conventionality on metaphor activation patterns

Broad contrasts comparing all types of metaphors to literal phrases do not account for two related facets of metaphor that have been the most extensively studied: familiarity and conventionality (Koller et al., 2022). The Career of Metaphor model posits that novel metaphors are initially understood as analogies, but after repeated encounters, the metaphor becomes conventionalized and acts as a categorical (i.e., a semantic) statement (Bowdle & Gentner, 2005; Gentner & Bowdle, 2008). The conventionalization of a metaphor can evolve to the point that its figurative nature is lost, becoming entirely lexicalized and indistinguishable from literal language (Schmidt et al., 2010). The graded salience hypothesis (Giora, 1997, 2003; Giora et al., 2000) proposes that when a metaphor becomes conventionalized, its figurative meaning becomes the most salient interpretation, and that salience—not figurativeness per se—determines the hemispheric laterality of semantic mapping during metaphor processing. The coarse semantic coding theory and graded salience hypothesis agree that literal language is predominantly processed within the LH while the RH is more adept at processing novel, figurative language. However, while the coarse semantic coding theory proposes that the RH plays a larger role than the left in processing conventional metaphors, the graded salience hypothesis attributes conventional metaphor processing to the LH.

Aligning more closely with the graded salience hypothesis, prior meta-analyses have indicated that only a small number of LH regions are activated to a greater extent for familiar (or conventional) metaphors over novel metaphors. Specifically, using effect size-signed differential mapping, Bohrn et al. (2012) found that familiar/conventional metaphors recruited LIFGorb (BA47), LMTG (BA 21), and left thalamus to a greater extent than novel metaphors. More recently, Huang et al. (2023) reported that a contrast activation likelihood estimation (ALE) analysis of conventional over novel metaphors yielded activated voxels within the left transverse temporal gyrus (BA 41) and the left insula. Analyses of novel over familiar/conventional metaphors present a somewhat more complicated story. Again, employing effect size-signed differential mapping, Bohrn et al. (2012) found greater activations for novel compared to familiar/conventional metaphors in the right insula extending to RIFGtri (BA 45) and right dorsal ACC (dACC, BA 32) but also in several LH regions, including peaks within LpITG (BA 37) extending into LpMTG (BA 21), LMFG (BA 46) spreading to LIFGop (BA 44), and the left parahippocampal cortex (BA 36). In Rapp et al. (2012), an ALE analysis of novel, non-salient over salient, nonliteral stimuli resulted in activations primarily within frontal LH and RH regions, including the bilateral middle frontal gyrus (MFG, BA 46) with peak activity within RMFG, LIFGtri (BA 45), RIFGorb (BA 47) extending into the right insula, and the right superior frontal gyrus (SFG; BA 8). Using independent ALE of novel over conventional metaphors, Huang et al. (2023) found activation within RIFGtri (BA 45) and the left medial frontal gyrus (BA 6), but their contrast ALE resulted in different activation patterns, specifically within bilateral SFG (left BA 6, right BA 8) and peak activation within LMFG (BA 46).

Despite some nuanced discrepancies between these studies, the collective meta-analytic findings suggest that compared to familiar/conventional stimuli, processing novel metaphors requires greater reliance on bilateral frontal regions, a finding that is not entirely explained by either the coarse semantic coding theory or the graded salience hypothesis. Moreover, while some of these frontal regions (IFG, mPFC) are often considered part of the core semantic network—implicated in controlled retrieval and selection processes—other regions (like the dACC, insula, SFG, and rostral MFG) are not (Hodgson et al., 2021; Jackson, 2021; Noonan et al., 2013). Instead, these regions are traditionally included in the multiple demand (MD) network, portions of the brain implicated in domain-general processes that exert top-down control over other networks when processing demands increase (Duncan, 2010; Fedorenko et al., 2011, 2013; Fedorenko & Thompson-Schill, 2014). In this vein, Schmidt and Seger (2009) argued that another variable driving processing of metaphors is task difficulty, with unfamiliar/unconventionalized metaphors being more difficult to process than conventionalized metaphors. While Schmidt and Seger’s (2009) arguments revolved around the role of the RH in processing difficult stimuli, instead it may be that activations within MD regions in *either* hemisphere should be observed for a contrast of novel over familiar/conventional metaphors if task difficulty is the primary driver in processing differences between these types of metaphors.

Another important consideration for the study of metaphor was summarized by Koller et al. (2022), who conducted a systematic review of the research methods used in past neuroimaging studies of figurative language and found vast inconsistencies between studies in the operational definitions and investigation of “familiarity” and “conventionality”. They found that the authors of 19 of 33 studies used “familiarity” and “conventionality” interchangeably, while the authors of three of the papers also used these terms synonymously with the term “salience”. The authors of 15 studies that equated familiarity to conventionality defined the construct according to subjective, individual-level familiarity reported by participants—a definition that Koller et al. (2022) argue aligns best with “familiarity” in the extant literature (Titone & Connine, 1994)—not according to the degree of entrenchment of phrases in everyday language, a definition that aligns more closely with “conventionality”. In other words, within a given culture, all conventional metaphors are highly familiar, but not all metaphors with some degree of familiarity are conventional. Relatedly, Koller et al. (2022) indicated that “novel” metaphors are often juxtaposed against both familiar and conventional metaphors—consistent with our summary above—yet, the majority of researchers (in 16 of 25 studies) defined “novel” stimuli as those with low participant-reported subjective familiarity, while other papers provided no operational definitions. Koller et al. (2022) also emphasize that while these variables are often measured along a continuum (via e.g., participant ratings from Likert scales), they are often binarized in neuroimaging analyses. Thus, a blurring of these terms has resulted in a confusing picture regarding the neural correlates of metaphor.

Using a clever design, Cardillo et al. (2012) circumvented these issues by conducting a study in which they manipulated the participants’ degree of familiarity with a set of completely novel metaphors prior to participants completing fMRI scanning. In this study, 120 metaphors were divided into three groups, two of which were shown to participants outside the scanner as part of a pre-exposure paradigm. Participants saw the first group of metaphors five times and the second group twice. The third group of metaphors were reserved as entirely novel, and participants did not see them until they completed the fMRI task. Cardillo and colleagues (2012) found that as the novelty of the metaphoric stimuli increased, so did the activation in bilateral IFGtri, LpMTG, and the right posterolateral occipital gyrus. LIFGtri and LpMTG are considered key hubs within the semantic control network (Jackson, 2021; Noonan et al., 2013), while the posterolateral occipital gyrus has been implicated in semantic integration when tasks require information being fed forward from visual regions to temporal cortex for semantic access (Binder et al., 2009). While Cardillo et al. (2012) provide important evidence regarding the Career of Metaphor, the findings cannot be used to disentangle effects of figurativeness from novelty/familiarity as all stimuli were metaphors and novel to varying degrees.

### 1.3. The current study

Thus, in this study, we extended the paradigm used by Cardillo et al. (2012) to include literal sentences in order to test hypotheses regarding how recruitment of LH semantic regions, RH homologues of those regions, and areas within the MD network vary as a function of sentence type (i.e., metaphorical versus literal) and novelty (i.e., completely novel versus somewhat familiar/pre-exposed). Neuroimaging data were collected using functional near-infrared spectroscopy (fNIRS), which is a flexible, relatively inexpensive technique capable of measuring cortical brain activation at a higher temporal resolution than fMRI. Thus, fNIRS may be useful for capturing potential differences in the temporal nature of LH versus RH recruitment during metaphor processing, but this method has been used in very few investigations of metaphor to date (i.e., only one paper out of 116 in Koller et al., 2022). Our research questions were as follows:

- *What are the differences in the neural correlates of a) metaphors versus literal sentences and b) novel metaphors versus familiar metaphors in healthy young adults?* Consistent with previous meta-analytic findings (Bohrn et al., 2012; Huang et al., 2023; Rapp et al., 2012; Yang, 2014), we hypothesized that metaphors would recruit core LH language regions (IFG, MTG, ITG, inferior parietal cortex), their RH homologues, and regions within the MD network not traditionally associated with language processing (bilateral MFG and SFG) to a greater extent than literal sentences across all participants. Consistent with Cardillo et al. (2012), we also predicted that activation within brain regions implicated in semantic control (i.e., bilateral IFGtri and the LpMTG) would be greater for novel relative to more familiar metaphors.
- *What are the associations between differences in task performance between conditions of interest (i.e., metaphors > literal sentences, novel > familiar metaphors) and brain activity?* Given that all metaphors included in this study were novel to some extent, we predicted that the difference in reaction times for metaphors versus literal sentences as well as novel versus more familiar metaphors would be associated with either activation within RH semantic regions—per the coarse semantic coding theory (Jung-Beeman, 2005) and graded salience hypothesis (Giora, 1997)—or within MD regions (Duncan, 2010; Fedorenko & Thompson-Schill, 2014) if task difficulty is the main driver of condition differences.

## 2. Materials and methods

### 2.1. Participants

To be eligible to participate, individuals needed to: 1) be an adult between the ages of 18 and 30 years, 2) have no history of neurological disease or major psychiatric disorders (e.g., schizophrenia, bipolar disorder, major depressive disorder), 3) have English as a primary language, 4) score above the education cutoff on the Mini Mental State Examination (Folstein et al., 1983), and 5) have normal or corrected-to-normal vision and hearing. Participants also needed to have at least 75% of fNIRS channels retained after preprocessing (see **2.5. fNIRS data processing**) for their data to be included in analyses. All 28 volunteers (14 women, 14 men^1^; 24 right-handed) met these criteria, participated in this experiment, and were compensated for their time. The sample size was established based on our goal of maximizing statistical power by approximately doubling the mean number of participants reported in a prior meta-analytic study of figurative language (*M* = 15.2, minimum = 6, maximum = 38; Bohrn et al., 2012).

The mean age of participants was 23.1 years (*SD* = 3.1, range = 18-29 years). Participants reported a mean number of 17.0 years of education (*SD* = 2.3, range = 13-22 years). Most participants identified their race as White (*n* = 11) or Asian (*n* = 14), with one participant indicating their race as “Biracial (White/Asian),” another as “Indian/Asian,” and a third as “Middle Eastern.” Participants separately reported their ethnicity as “non-Hispanic” (*n* = 23), White (*n* = 1), Caucasian (*n* = 1), non-Hispanic Indian (*n* = 1), and Persian/Iranian (*n* = 1); one individual chose not to disclose their ethnicity. Four participants were monolingual English speakers; 22 were bilingual/multilingual; and two individuals did not specify their language history. Twenty-three individuals reported their first language as English (*n* = 20) or English plus another language (*n* = 3). Other reported first languages included Hindi (*n* = 3), Marathi (*n* = 2), Tamil (*n* = 2), Gujarati (*n* = 1), and Russian (*n* = 1), with two participants reporting more than two primary languages. All participants reported being fluent in English by age seven, a requirement for inclusion in the study and qualified by their report of consistent use of English in a home and/or school setting by that age. Our rationale for including bi/multilingual individuals was to achieve a more diverse sample that is more representative of the growing global majority than past fNIRS language research (Girolamo et al., 2023; Kwasa et al., 2023). Subject-level demographic data are reported in Supplemental Table 1.

All participants provided written consent in accordance with the policies of the Declaration of Helsinki of 1975 and the procedures approved by the Institutional Review Board at Northeastern University under protocol #21-06-11.

### 2.2. Experimental stimuli

As previously referenced, this study was based on and extends an fMRI study conducted by Cardillo et al. (2012). A set of 120 sentences (60 literal, 60 metaphorical) were selected from the stimuli normed by (Cardillo and colleagues (2010). Within the Cardillo et al. (2010) final set, each sentence has a base term, such as “flush.” In the current study, each base term was used in at least two sentences, one metaphorical (e.g., *His memoirs were a toilet flush*) and one literal (e.g., *The only noise was a toilet flush*). Half of the phrases were predicate phrases in which the base term was a verb (e.g., *The panther growled at the photographer* or *the cruise ship growled at the fishing boat*), and half were nominal phrases in which the base term was a noun (e.g., *The puppy’s grasp was a firm tug* or *The shop display was a gentle tug*). Predicate sentences were in the past tense, in the form of NOUN VERB-ED PREPOSITION INDIRECT-OBJECT (e.g., *The football team lurched through the season*). Nominal phrases were also in the past tense, using either the singular or plural form of the verb to be, in the form of NOUN TO-BE NOUN (e.g., *The winter was a heartbroken limp*). To account for potential modality-specific activations evoked by the target base term, half of the phases included an auditory-related base term (e.g., “flush”, “whistle”, “giggle”), and the other half included a motion-related base term (e.g., “jog”, “dig”, “crawl”). To maintain balance across sentence types (i.e., auditory versus motion, nominal versus predicate), five base terms were used four times, resulting in a total of 56 base terms in our experimental stimuli.

Metaphorical and literal sentences in our experiment were matched on length in terms of number of words (*t*_118_ = -0.42, *p* = 0.675; mean length: metaphorical = 6.40 words, literal = 6.35 words) and letters (*t*_118_ = -1.25, *p* = 0.212; mean length: metaphorical = 29.15 letters, literal = 27.97 letters). We also compared metaphorical and literal sentences from our stimulus subset on Likert ratings of imageability, naturalness, familiarity, and figurativeness, using norms from Cardillo et al. (2010). Consistent with findings from Cardillo et al. (2010) on the full dataset, our stimuli differed in expected ways: literal sentences had significantly higher imageability (*t*_114_ = 18.59, *p* < 0.001), naturalness (*t*_112_ = 20.63, *p* < 0.001), and familiarity (*t*_114_ = 19.39, *p* < 0.001) ratings than metaphorical sentences whereas literal sentences had significantly lower figurativeness ratings (*t*_118_ = -31.79, *p* < 0.001) than metaphorical sentences. See Supplemental Table 2 for the full phrase list used in the current study.

### 2.3. Behavioral data acquisition

Following completion of the consent and intake paperwork, participants completed a battery of behavioral assessments as part of the larger scope investigation of the neural bases of cognitive-linguistic processes in adults with and without a history of stroke. While examining relationships between offline behavioral task performance and fNIRS brain activity is outside the scope of this study, data used to characterize the sample are reported at the single-subject level in Supplemental Table 1.

#### 2.3.1. Familiarization task

Active, elaborative processing has been shown to have a strong effect on retention (Benjamin & Bjork, 2000). Thus, like Cardillo et al. (2012), prior to the fNIRS experiment, the participants were asked to engage in active processing of the metaphors and literal sentences, including making judgments about their characteristics (i.e., ease of comprehension, imageability, figurativeness). Cardillo et al. (2012) split their stimuli into three sets: 1) high exposure (i.e., sentences presented five times to participants prior to the fMRI experiment), 2) moderate exposure (i.e., sentences presented twice to participants prior to the fMRI experiment), and 3) no exposure/novel (i.e., sentences that were not pre-exposed prior to the fMRI experiment). In our study’s pre-exposure (familiarization) phase, our stimuli were split evenly between sentences that participants were exposed to during the familiarization task (i.e., 50% of metaphorical sentences with matched literal pairs) and sentences that were excluded from the familiarization task and seen for the first time in the fNIRS experiment. The reasons for deviating from Cardillo et al. (2012)’s three-set system were to reduce the length of the experiment (as our unique inclusion of literal sentences doubled the number of sentence stimuli) and simplify the fNIRS comparison contrasts. Sentences were assigned to the pre-exposure/familiar or no-exposure/novel conditions randomly and were counterbalanced so that approximately equal numbers of each sentence type (i.e., auditory versus motion, nominal versus predicate) were included in the novel and familiar subsets. This familiarization experiment was split into two runs of 30 sentences each, split evenly between metaphorical and literal trials and presented in PsychoPy version 3.2.10 (Peirce et al., 2019). During the familiarization task, the participant was presented with each sentence three times and asked to answer one of three questions regarding imageability, figurativeness, or comprehensibility of sentences using a 7-point Likert scale response. Items were presented in fully randomized order. The three prompts were:

1. *Imageability*: Please rate how quickly and easily each sentence brought a visual image to mind using a scale of 1 (very imaginable) to 7 (very unimaginable) by clicking on the sliding scale below with your mouse.
2. *Figurativeness*: Please rate how literal of an interpretation is suggested by this sentence using a scale of 1 (very literal) to 7 (very figurative) by clicking on the sliding scale below with your mouse.
3. *Comprehensibility*: Please rate how easy or hard it is to make sense of this phrase using a scale of 1 (very easy) to 7 (very hard) by clicking on the sliding scale below with your mouse.

#### 2.3.2. fNIRS task

The fNIRS experiment was split into two runs, with a total of 120 phrases split into four conditions: 1) familiar literal (FAM/LIT, *n* = 30 sentences), 2) novel literal (NOV/LIT, *n* = 30 sentences), 3) familiar metaphorical (FAM/MET, *n* = 30 sentences), and 4) novel metaphorical (NOV/MET, *n* = 30 sentences). Each run was 7 minutes and 16 seconds in duration. The experiment followed a block design, with each block containing five sentences from a given condition. The duration of each sentence trial was five seconds, resulting in 25s blocks. Each run of the experiment contained a total of 12 blocks (three blocks of each condition) presented in a randomized order. An inter-block-interval with a randomized length ranging from 5 to 12 seconds (*M* = 10s) was presented between blocks during which the participants were asked to rest and fixate on a cross presented in the middle of the screen. See Figure 1 for an example of the task time series.

**Figure 1.**
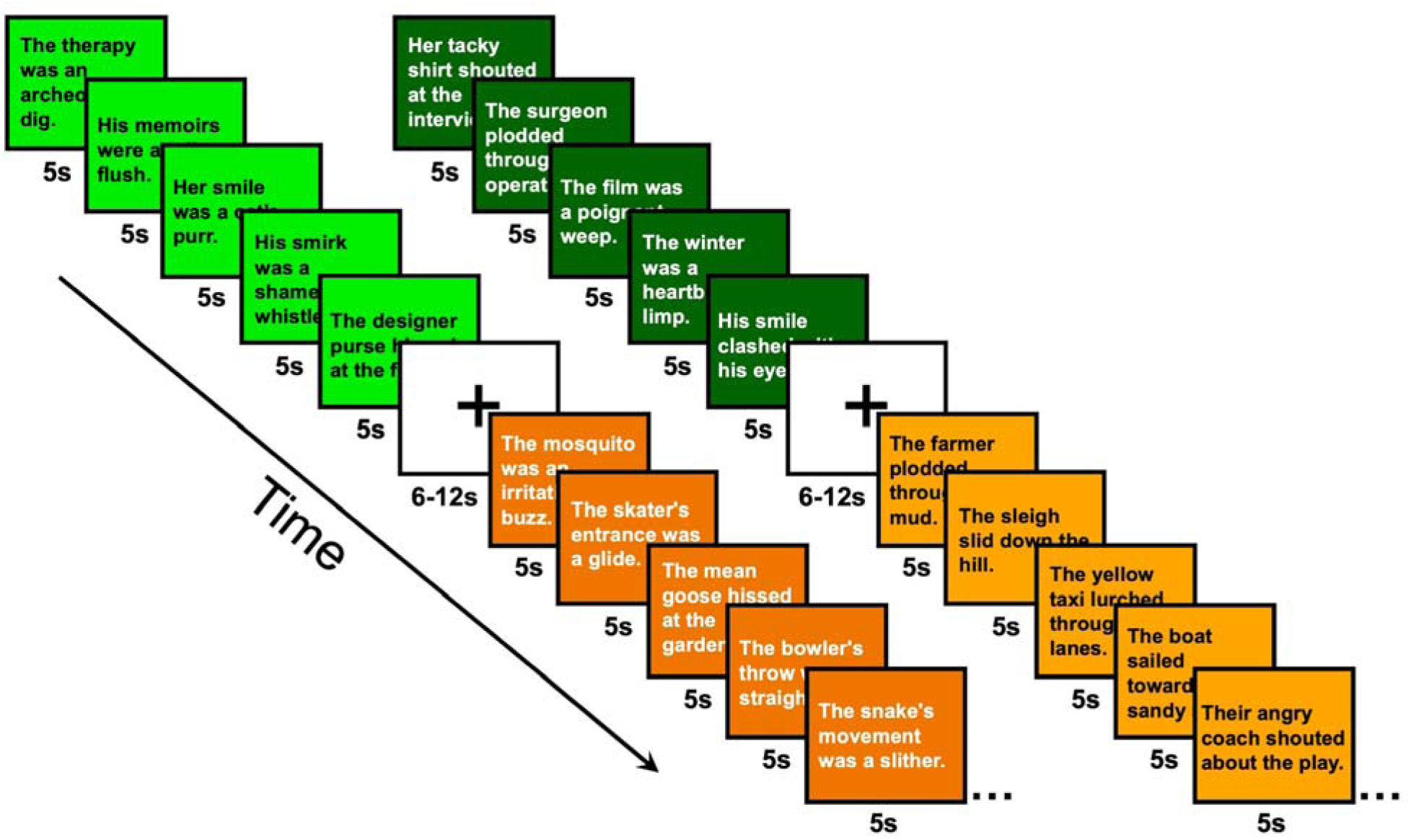
Example time series of the fNIRS task. The time series included 25s blocks containing either five familiar literal (light orange), novel literal (dark orange), familiar metaphorical (neon green), or novel metaphorical (dark green) sentences per block. Each block was separated by a jittered 6-12s rest period.

For each trial, participants were asked to provide a button press response for each sentence to encourage active processing during the task. Participants saw the following instructions at the beginning of each run: “You will see a sentence. After reading each sentence, please rate how easy or difficult it is to make sense of the phrase you are reading. Use the buttons to indicate your response, where GREEN is easy and RED is difficult.” For this judgment, two outcome measures were recorded: 1) the reaction time (RT) between the participant being presented with the stimulus (in seconds) and 2) which key they selected as their response (0 = easy, 1 = hard).

#### 2.3.3. Base term task

After completing both fNIRS runs, each participant completed a brief PsychoPy experiment in which they were presented with each base term (*n* = 56) and asked to indicate (via button press) whether they understood the meaning of the term (i.e., if they could use it correctly in a sentence) or not. The purpose of this task was to ascertain if any base terms were unfamiliar to the participant, which could impact their results. Participants reached near-ceiling performance on this task (*M* = 95.28% “yes” responses), suggesting that unfamiliarity with vocabulary was not a driving force in performance.

### 2.4. fNIRS data acquisition

In fNIRS imaging, low-powered light-emitting diodes (called sources) arranged across the scalp surface project near-infrared optical signals through the scalp, skull, and meningeal layers into the cortical surface (typically at a depth of 1-2 cm). Oxygenated and deoxygenated hemoglobin chromophores in tissues within the optical path absorb incoming signals, and the quantity of photons that return to the scalp surface are measured by photodiode detectors situated nearby a paired source. Hemodynamic responses are obtained by converting relative changes in chromophore intensity to changes in optical density, followed by conversion of changes in optical density to relative changes in oxygenated and deoxygenated hemoglobin concentrations via the modified Beer-Lambert Law (Kocsis et al., 2006). Thus, similar to fMRI, fNIRS provides an indirect measure of brain activity through specification of the hemodynamic response function to task stimuli.

In this study, neuroimaging data were collected using two continuous wave 8x8 NIRx NIRSport2 devices (https://nirx.net/nirsport) at a sampling rate of 11.2 Hz. The devices were daisy chained together during data acquisition, effectively creating an array of 16 high-powered, dual-tipped LED sources and 16 dual-tipped silicon photodiode detectors. During data acquisition, sources projected near-infrared light at two wavelengths, 760 and 850 nm. Each paired source and detector that were spaced ≤ 30 mm apart formed a long-distance measurement channel, resulting in 44 long-distance channels arranged over perisylvian areas and evenly split between cerebral hemispheres. One detector was used to acquire data from eight short-distance channels that were situated 8mm from each paired source and distributed across the montage, with two posterior and two anterior short-distance channels in each hemisphere. Short-distance channels were used to capture physiological signals of non-interest (e.g., intra-arachnoid blood flow) that were regressed out in data processing.

The source-detector probe was created to prioritize capturing data from regions typically associated with language processing in the LH, their RH homologues, as well as portions of the bilateral pre-frontal cortices implicated in executive processes. Regions covered by the probe included: 1) portions of the dorsolateral PFC, rostral lateral PFC, and frontal pole which included MFG and anterior SFG, 2) IFGtri and IFGop, 3) the precentral gyrus, 4) mid-to-posterior STG, 5) mid-to-posterior MTG, 6) posterior ITG, 7) angular gyrus (AG) and 8) supramarginal gyrus (SMG). AtlasViewer (Aasted et al., 2015) was used to obtain anatomical labels for each channel by registering the probe to the head surface and projecting data onto the Colin brain atlas (Collins et al., 1998). Following this registration, AtlasViewer assigned a Montreal Neurological Institute (MNI) coordinate and corresponding anatomical label according to the Automated Anatomical Labeling (AAL) Atlas (Tzourio-Mazoyer et al., 2002) for each channel based on the midpoint of each source-detector pair. Using an approach utilized in past fNIRS research (Gilmore et al., 2021; Li et al., 2020), channels were then grouped according to regions of interest (ROIs) using the average MNI coordinate location. See Figure 2 for a visualization of the probe design and sensitivity profile and Table 1 for the list of ROIs by channel and MNI coordinates.

**Figure 2.**
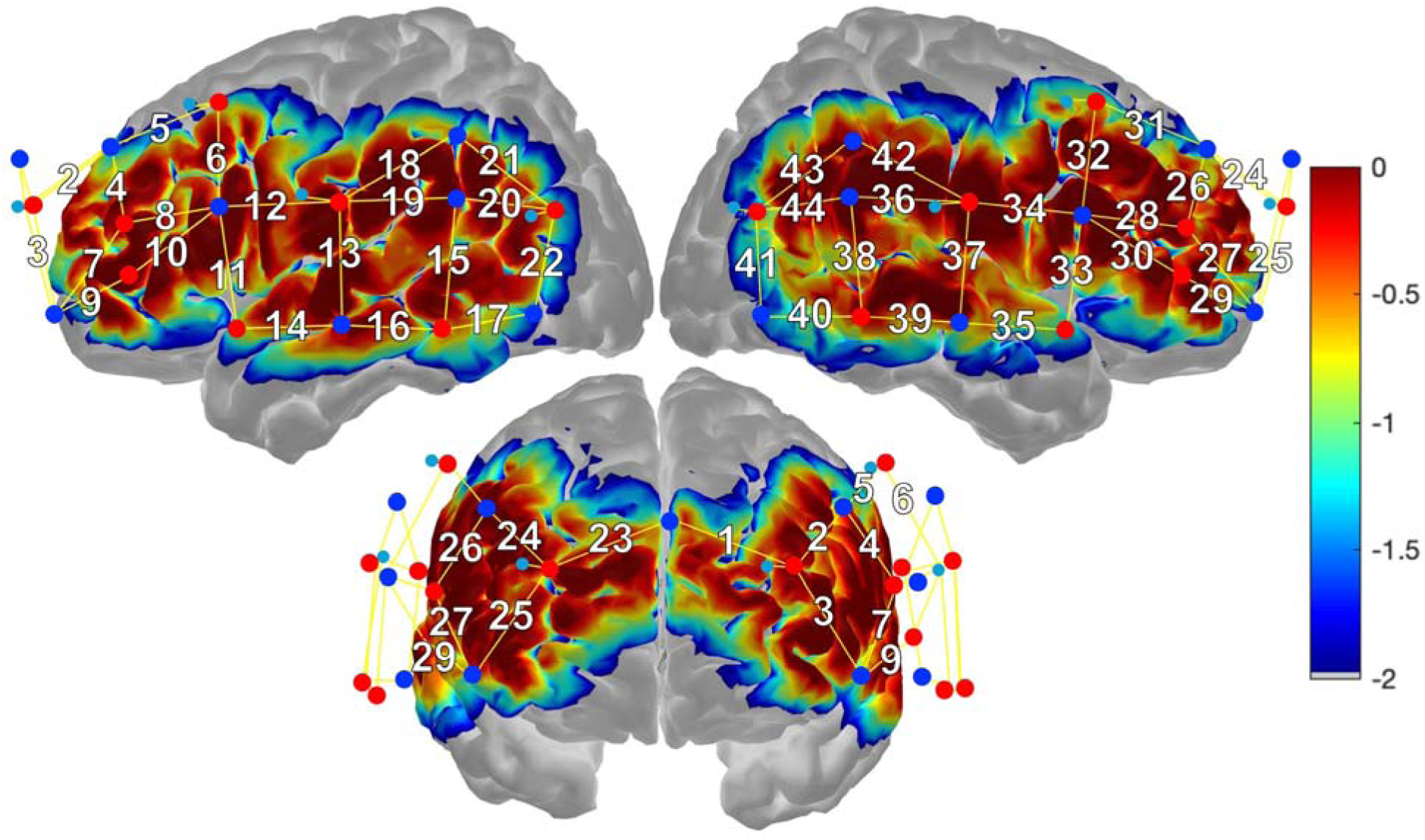
fNIRS probe and sensitivity profile. Sources (red circles) and detectors (blue circles) arranged over perisylvian areas form 44 long-distance channels (solid yellow lines, white numbers). Short channels are formed by detectors (light blue circles). The sensitivity profile (right) depicts areas of higher (warm colors) and lower (cool colors) measurement sensitivity.

**Table 1.** Regions of Interest (ROIs) by Channel and MNI Coordinates.

| ROI | Left Hemisphere | Right Hemisphere |
| --- | --- | --- |
| Superior Frontal Gyrus (SFG) | Ch1 (-10 55 22);<br>Ch2 (-22 46 21) | Ch23 (16 69 28) |
| Middle Frontal Gyrus (MFG) | Ch3 (-30 59 6);<br>Ch4 (-38 37 20) | Ch24 (33 57 28);<br>Ch25 (40 67 9);<br>Ch26 (42 32 17) |
| Inferior Frontal Gyrus, pars triangularis (IFGtri) | Ch5 (-33 22 24);<br>Ch6 (-40 16 25);<br>Ch7 (-40 49 6);<br>Ch8 (-35 22 22);<br>Ch9 (-39 47 3);<br>Ch10 (-45 28 19);<br>Ch11 (-55 13 13) | Ch27 (44 46 2);<br>Ch28 (46 24 19);<br>Ch29 (55 53 1);<br>Ch30 (56 27 16) |
| Inferior Frontal Gyrus, pars opercularis (IFGop) | Ch12 (-48 7 25) | Ch31 (45 26 32);<br>Ch32 (55 17 31) |
| Superior Temporal Gyrus (STG) | Ch13 (-53 -14 9) | Ch36 (63 -26 23);<br>Ch37 (65 -11 9);<br>Ch38 (72 -43 12) |
| Middle Temporal Gyrus (MTG) | Ch14 (-62 -1 -6);<br>Ch15 (-43 -38 8);<br>Ch16 (-49 -26 -6) | Ch39 (53 -26 -8);<br>Ch40 (58 -46 -5);<br>Ch41 (43 -56 12) |
| Inferior Temporal Gyrus (ITG) | Ch17 (-47 -46 -6) | N/A |
| Supramarginal Gyrus (SMG) | Ch18 (-56 -24 35);<br>Ch19 (-46 -24 25);<br>Ch20 (-56 -52 26) | Ch42 (50 -23 32) |
| Angular Gyrus (AG) | Ch21 (-53 -53 32) | Ch43 (45 -46 30);<br>Ch44 (59 -50 25) |

Notes: Channels (Ch) are labeled with the number that corresponds labels in Figure 2. Hemispheric asymmetry resulted in homologous channels being assigned to different regions of interest. Most notably, this resulted in the most posterior-inferior channel in the left hemisphere to be assigned to the left ITG whereas the homologous channel was assigned to the right MTG. Montreal Neurological Institute (MNI) coordinates are listed according to x-, y-, and z- axis locations. Abbreviations: N/A = not applicable.

At the beginning of the session, measurements of each participant’s head circumference, nasion to inion distance, and distance between the left and right pre-auricular points were obtained. The optodes were transferred to and stabilized within an appropriately sized 10-20 EASYCap (EASYCap GmbH, Woerthsee-Etterschlag, Germany) to ensure proper placement of source-detector optodes relative to the participants’ anatomical markers (e.g., nasion, inion, and left and right pre-auricular points) and to each other (i.e., ≤ 30 mm spacing). Research assistants placed the fNIRS cap on the participant’s head and adjusted the position of the cap relative to fiducials (e.g., Cz on the cap) and anatomical landmarks (i.e., nasion, inion, and pre-auricular points). Once the cap was donned, photos were taken of the participant from the front, back, and each side as a quality check to review and verify placement later. The position of the optodes was assessed manually to ensure good contact with the participant’s scalp, and then the fNIRS cap was covered with an opaque shower cap intended to reduce external light interference.

fNIRS data were recorded using NIRx Aurora software (https://nirx.net/software). Prior to the start of the fNIRS experiment, the signal optimization protocol in Aurora was run to assess the global signal, the amount of dark noise (reflecting the detectors’ capacity to detect incoming light), and the coefficient of variation in the signal (reflecting the signal to noise ratio). If any readings were suboptimal (e.g., critical coefficient of variation in Aurora at > 7.5%), the appropriate adjustments—such as gently moving hair aside with a wooden dowel or switching to higher tension optode tops to obtain better optode-scalp contact—were made to improve readings such that at least 75% of channels exhibited excellent or acceptable signal levels. Signals were monitored throughout data acquisition, and signal optimization and cap/optode adjustments were re-run as needed between task runs.

### 2.5. fNIRS data processing

Task data were preprocessed in Homer3 (Huppert et al., 2009) twice, first by replacing the default four condition markers (i.e., FAM/LIT, NOV/LIT, FAM/MET, NOV/MET) with markers that reflected the broader contrast of interest (i.e., LIT [literal] versus MET [metaphorical]) and second by retaining the four default markers to evaluate the effect of novelty on metaphorical processing (i.e., FAM/MET versus NOV/MET). The same preprocessing steps were followed in each analysis. First, channels with SNR < 5.0 were pruned, and raw signals were converted to changes in optical density (OD). Motion correction was performed using the splinesg routine, a combination of spline interpolation and Savitzky-Golay filtering (Jahani et al., 2018). An artifact detection step was subsequently run to flag lingering abrupt shifts (over 200 ms) in the OD data, and stimulus markers surrounded by uncorrected artifacts (-5s to 10s) were rejected, resulting in the removal of the corresponding block. The OD data were then bandpass filtered with a low-pass filter of 0.5 Hz to remove high-frequency noise (e.g., cardiac pulsations) and a high-pass filter of 0.01 Hz to remove low-frequency noise (e.g., slow signal drift). Following that step, changes in OD were transformed into changes in oxyhemoglobin (HbO) and deoxyhemoglobin (HbR) concentrations via the modified Beer-Lambert Law (Kocsis et al., 2006) using a partial path length factor of 1.0. Finally, changes in HbO and HbR concentrations for each condition were calculated using an ordinary least squares general linear model (GLM) with the HRF modeled as a sequence of consecutive Gaussians (Diamond et al., 2006) and the time range set to 2s before to 30s after the onset of the first trial of each block. Within the GLM, physiological signals of non-interest were regressed out using the data from the short-distance channel data mostly highly correlated with data from a particular long-distance channel (Gagnon et al., 2012; Yücel et al., 2015).

AtlasViewer (Aasted et al., 2015) was used to project HbO concentrations onto the brain in stereotactic space for each condition of interest. Image reconstruction was performed with the short separation channel length set to 8mm; other default settings (i.e., α regularization threshold of 0.01) were preserved. Changes in HbO for each condition of interest (i.e., MET, LIT, NOV/MET, and FAM/MET) were assessed in 5s epochs within the 0-30s time range.

### 2.6. Statistical analyses

Statistical analyses were conducted in R (R Core Team, 2024). To determine statistical differences in brain activity for each contrast of interest (i.e., MET > LIT and NOV/MET > FAM/MET) for aim #1, each participant’s HbO data were first downsampled by averaging the 7-8 datapoints collected within each 1s time window^2^ over the analysis time range (0-30s) for each condition. These data were then averaged across channels within each ROI per the channel assignments in Table 1. To determine the differences in activity between metaphorical and literal sentences as well as between novel and familiar metaphors, we ran a series of paired sample t-tests for each 1s time window for each ROI using the ‘rstatix’ package (Kassambara, 2023). Correction for multiple comparisons was performed at the false discovery rate (FDR, *q* < 0.05) across all time windows and ROIs for the two contrasts of interest (i.e., MET > LIT and NOV/MET > FAM/MET).

To address aim #2, we first examined how the reaction times (RTs) and key responses from the sentence comprehensibility judgments during the fNIRS experiment varied by condition via mixed effects models using the ‘lme4’ (Bates et al., 2015) and ‘lmerTest’ (Kuznetsova et al., 2017) packages. We first specified a base linear (for RT) and logistic (for key response) mixed effects model that included random intercepts for each participant and item. Using Analyses of Variance (ANOVAs), the base model was compared to the model of interest, which included a fixed effect of condition (i.e., FAM/LIT, NOV/LIT, FAM/MET, and NOV/MET) in which the reference level was set to FAM/LIT. Pairwise comparisons between conditions for significant fixed effects were evaluated using the ‘emmeans’ package (Lenth, 2024) with Tukey adjustment.

To answer our main question within aim #2, we completed additional post-processing of the behavioral data to generate variables that reflected the behavioral and neural differences of contrasts of interest (i.e., MET > LIT and NOV/MET > FAM/MET). The behavioral effect of sentence type was quantified by subtracting the averaged RT and percentage of “hard” key responses (indicating the sentence was challenging to comprehend) for the LIT condition from RTs/key responses for MET condition for every participant. Similarly, the behavioral effect of novelty was quantified by subtracting the average RT/percent “hard” key responses for FAM/MET from the average RT/percent “hard” key responses for NOV/MET for each individual. For the fNIRS data, we specified two sets of Bayesian linear mixed effects models (one for each contrast) using the ‘blme’ package (Chung et al., 2013) with the ‘optimx’ optimizer (Nash & Varadhan, 2011, 2011). Within each model, the dependent variable was the downsampled HbO time series data for a given ROI with a fixed effect of the interaction of condition by time, and by-participant random intercepts and slopes for time, condition, and their interaction. The inclusion of this random effect allowed for each participant to have their own baseline and unique pattern of change in HbO over time across conditions, thereby yielding subject-specific slopes that reflected that individual’s difference in activation patterns between conditions of interest for each ROI. Subject-specific slopes were extracted, standardized, and used in subsequent regression analyses.

To evaluate brain-behavior relationships, we used two-tailed Least Absolute Shrinkage and Selection Operator (LASSO) regression (Tibshirani, 1996) within the ‘selective Inference’ package (Tibshirani et al., 2019). LASSO regression is ideal for neuroimaging analyses because it can account for multicollinearity between variables as well as a large number of predictors relative to the sample size by imposing a shrinkage parameter and variable selection process that results in a sparse, optimal model (Meinshausen & Yu, 2009). We used standard features, including leave-one-out cross-validation to select a LASSO penalty parameter (λ) that resulted in the minimum mean cross-validated error. In each model, the dependent variable was the RT or key response difference metric, and the independent variables were the single-subject slopes for each ROI.

### 2.7. Data availability statement

The datasets associated with study are available within the Open Science Framework (Meier & Schwartz, 2025).

## 3. Results

An average of 1.6% (*SD* = 3.3%) of the long-distance channels were pruned across the sample. Nineteen participants (out of 28 participants) had no pruned channels, whereas the participant with the noisiest data had six pruned channels (out of 44). For aim #1, data from retained channels were condensed to the ROIs, and missing values from pruned channels were removed from analysis. See Supplemental Table 3 for a list of pruned channels for each subject by ROI. LASSO regression fails with missing values; therefore, to maximize statistical power for brain-behavior analyses in aim #2, missing data from pruned channels were imputed via predictive mean matching (with five imputations x50 iterations) using the ‘mice’ package (Buuren & Groothuis-Oudshoorn, 2011).

### 3.1. Aim #1: Differences in brain activity between conditions of interest

#### 3.1.1. Metaphorical versus literal sentences

Figure 3 shows the image reconstructions of HbO concentrations for MET (metaphorical sentences; Figure 3A) and LIT (literal sentences; Figure 3B) conditions over the 0-30s time block. Increases in HbO peaked within the 10-20s range for both conditions across most activated ROIs. As expected, increases in HbO concentrations were greater in the left than right hemisphere for each time window for both conditions. Indeed, the time series plots (see Figure 4) indicate that the only RH ROI that was significantly activated for both conditions was RIFGtri, indicated by the characteristic curve of the hemodynamic response in this region (i.e., a rise from the 0 baseline to a peak around 10-20s, followed by a decrease back to baseline). As shown in image reconstruction (Figure 3) and confirmed with the time series plots (Figure 4), left hemisphere regions significantly activated by both conditions included LIFGtri, LIFGop, LSTG, LMTG, and LITG. Bilateral SFG was not significantly recruited for either condition, nor were RMFG, RSTG, or RSMG.

**Figure 3.**
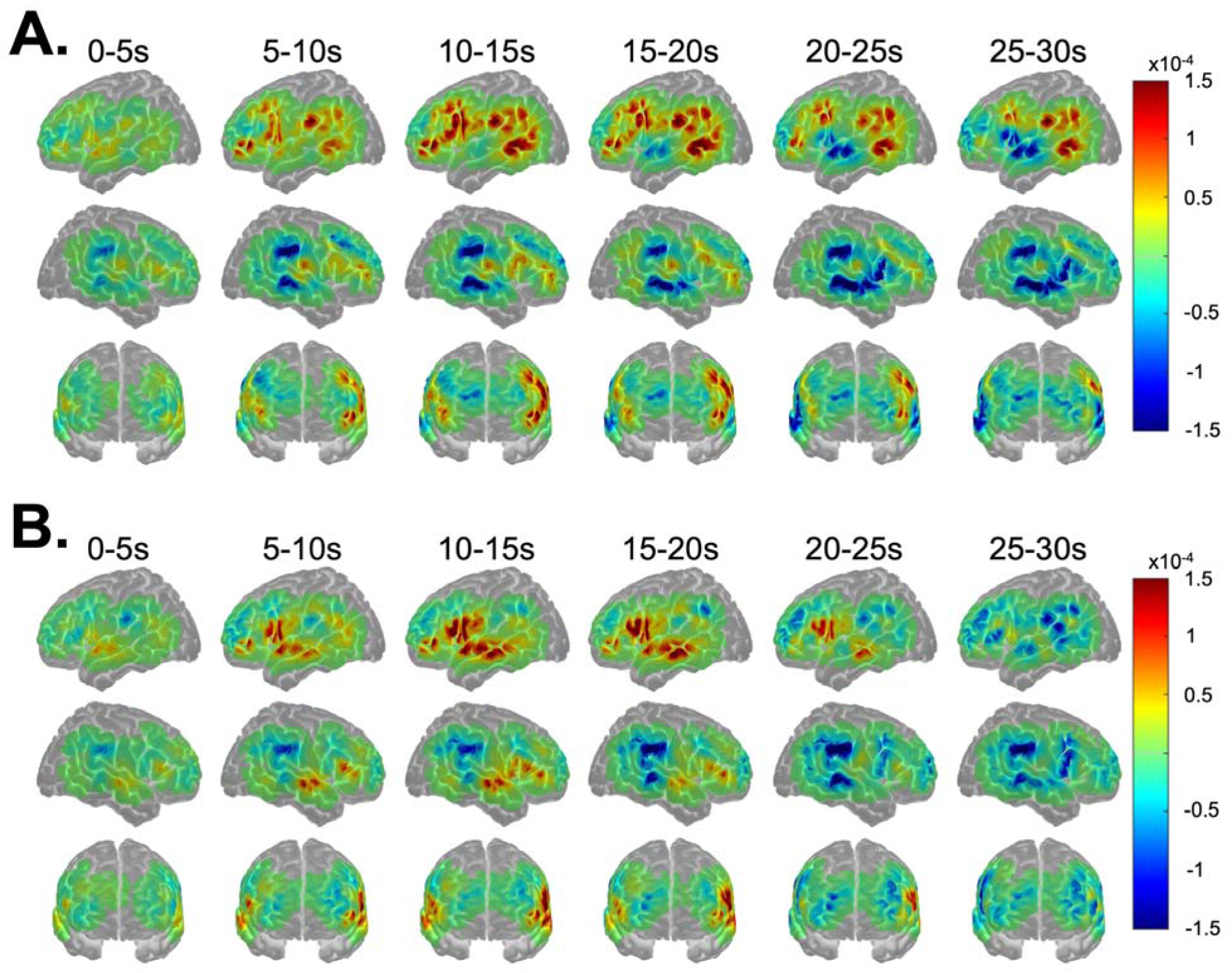
Image reconstruction of changes in oxyhemoglobin (HbO) for A) metaphorical and B) literal sentences. Warm and cool colors reflect, respectively, increases and decreases in HbO over the time block in 5s increments.

**Figure 4.**
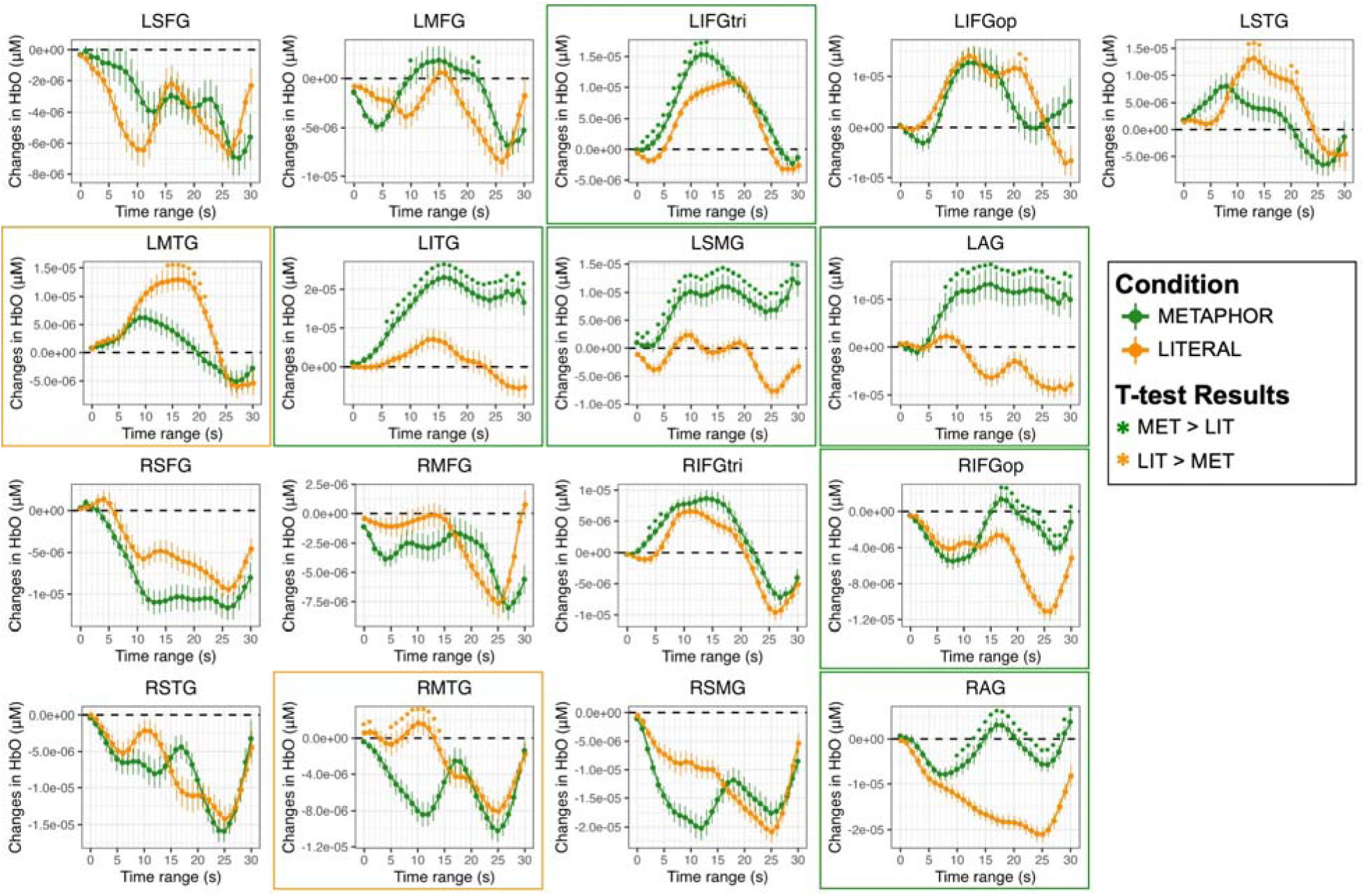
Comparison of changes in oxyhemoglobin (HbO) between metaphorical (MET) and literal (LIT) sentences for each region of interest (ROI). Significant differences between task conditions are based on paired t-tests, corrected for multiple comparisons across 1s epochs and ROIs (\**q* < 0.05), and denoted by asterisks in the color that reflects the direction of the effect, where green indicates greater changes in HbO for MET > LIT and orange indicates greater HbO changes for LIT > MET. Borders denote ROIs for which a characteristic increase in HbO was observed and the between-condition difference persisted for at least five seconds. Abbreviations: L = left; R = right; SFG = superior frontal gyrus; MFG = middle frontal gyrus; IFGtri = inferior frontal gyrus, pars triangularis; IFGop = inferior frontal gyrus, pars opercularis; STG = superior temporal gyrus; MTG = middle temporal gyrus; ITG = inferior temporal gyrus; SMG = supramarginal gyrus; AG = angular gyrus.

Left-lateralized brain activity was significantly greater for the MET than LIT condition across several ROIs, including LMFG over a brief window (10s and 21-22s, *q* < 0.05), LIFGtri (from 1-13s, *q* < 0.05), LITG (from 6-30s, *q* < 0.05), LSMG (from 0-2s and 4-30s, *q* < 0.05), and LAG (from 8-30s, *q* < 0.05). Despite the short duration and weak magnitude of the effects in most RH ROIs, greater increases in HbO for MET than LIT were noted in RIFGop (from 17-30s, *q* < 0.05) and RAG (from 10-30s, *q* < 0.05). There was also a numerical trend of greater increases in HbO in RIFGtri across the entire time series for MET compared to LIT, but most of the comparisons did not survive correction for multiple tests except for a brief period from 4-6s (*q* < 0.05).

The only LH ROI that demonstrated a consistently greater response for the LIT than MET condition was LMTG (from 14-21s, *q* < 0.05), but LSTG also exhibited a couple brief periods of significantly greater change in HbO for LIT compared to MET (from 12-14s and 20-21s, *q* < 0.05) A brief increase in HbO in RMTG for the LIT condition was also significantly higher than RMTG activity observed for the MET condition (from 0-1s and 5-14s, *q* < 0.05). The complete t-test outputs of the comparisons between MET and LIT conditions for each second and ROI are reported in Supplemental Table 4.

#### 3.1.2. Novel versus familiar metaphors

Figure 5 shows the image reconstructions of HbO concentrations for the NOV/MET (novel metaphors; Figure 5A) and FAM/MET (familiar metaphors; Figure 5B) conditions. As expected based on the activation patterns for the broader MET condition, increases in HbO over the time block were highly left lateralized for both NOV/MET and FAM/MET. Peak activation was again noted in the 10-20s range for both conditions. However, the onset of increases in HbO in most LH ROIs for NOV/MET lagged activation onsets for FAM/MET by around five seconds, with little increases in HbO for NOV/MET in the 5-10s window (except for in LMFG) compared to widespread increases in HbO for FAM/MET in that same time range. As visualized in the image reconstructions (Figure 5) and supported by the time series plots (see Figure 6), regions significantly activated by both NOV/MET and FAM/MET included LIFGtri, LIFGop, LSTG (albeit briefly and lacking the canonical curve), LITG, LSMG, LAG, and RIFGtri. Bilateral SFG, RSTG, and RSMG were not significantly activated for either condition.

**Figure 5.**
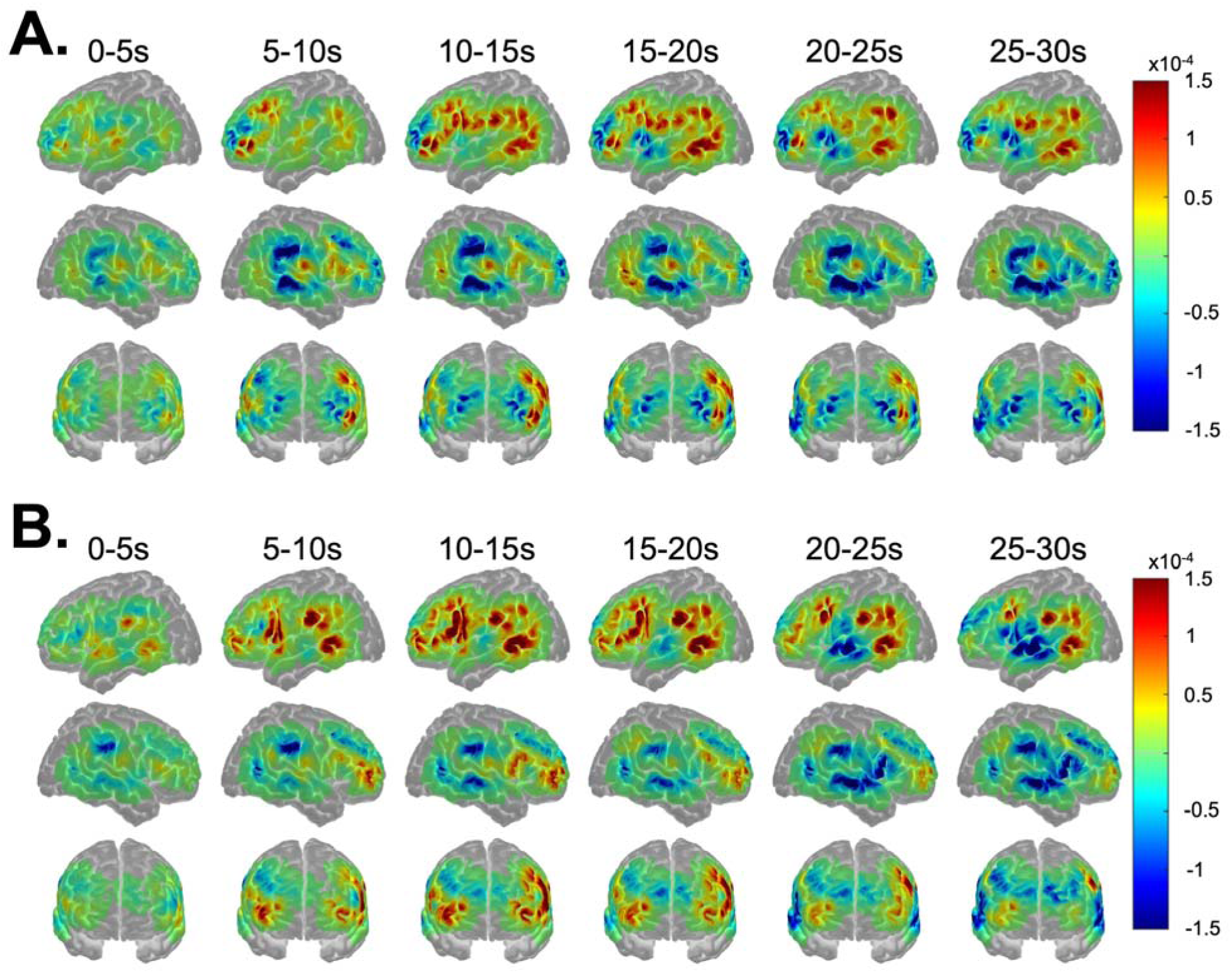
Image reconstruction of changes in oxyhemoglobin (HbO) for A) novel and B) familiar metaphorical sentences. Warm and cool colors reflect, respectively, increases and decreases in HbO over the time block in 5s increments.

**Figure 6.**
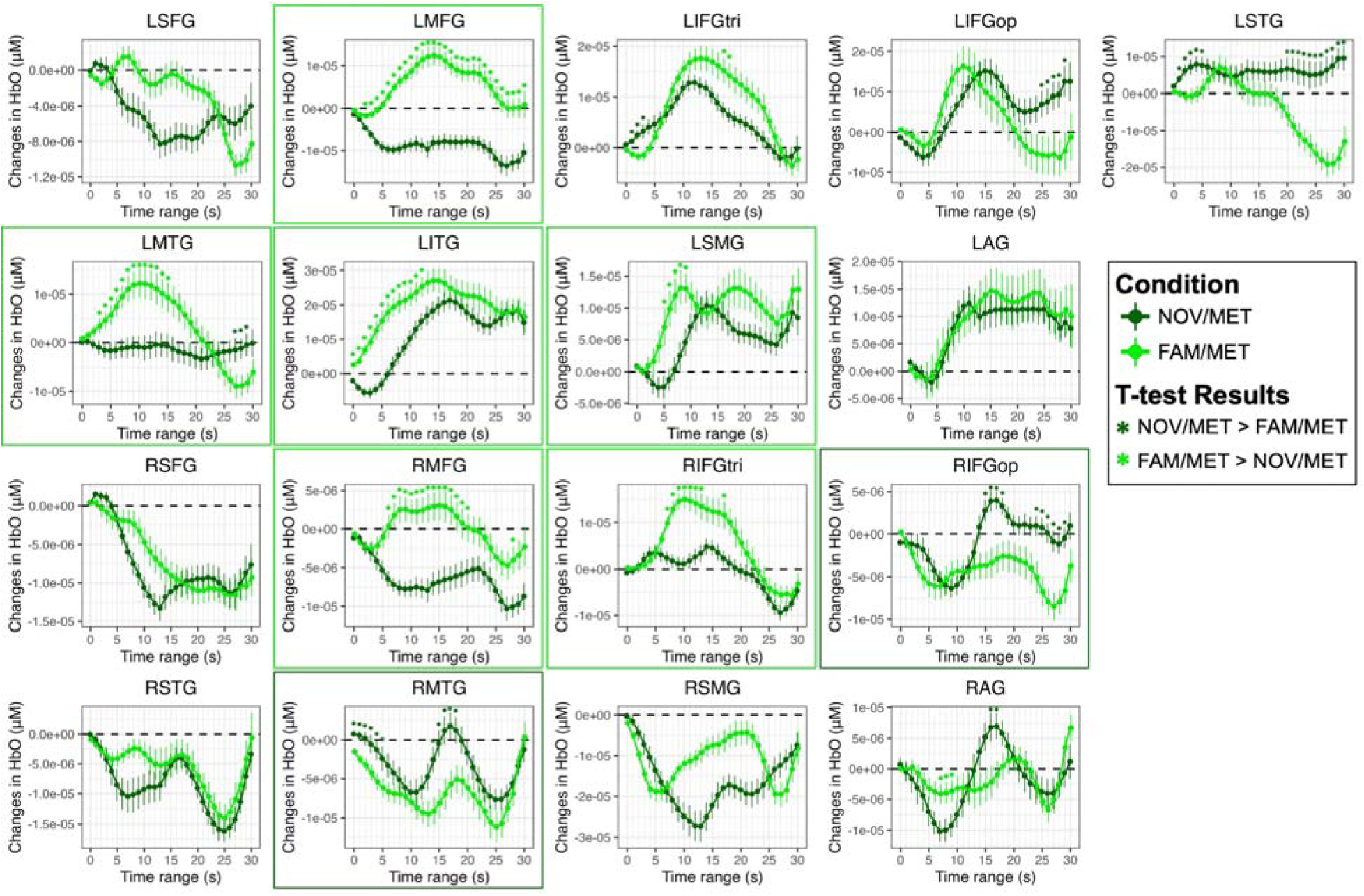
Comparison of changes in oxyhemoglobin (HbO) between novel metaphorical (NOV/MET) and familiar metaphorical (FAM/MET) sentences for each region of interest (ROI). Significant differences between task conditions are based on paired t-tests, corrected for multiple comparisons across 1s epochs and ROIs (\**q* < 0.05), and denoted by asterisks in the color that reflects the direction of the effect, where dark green indicates greater changes in HbO for NOV/MET > FAM/MET and neon green indicates greater HbO changes for FAM/MET > NOV/MET. Borders denote ROIs for which a characteristic increase in HbO was observed and the between-condition difference persisted for at least five seconds. Abbreviations: L = left; R = right; SFG = superior frontal gyrus; MFG = middle frontal gyrus; IFGtri = inferior frontal gyrus, pars triangularis; IFGop = inferior frontal gyrus, pars opercularis; STG = superior temporal gyrus; MTG = middle temporal gyrus; ITG = inferior temporal gyrus; SMG = supramarginal gyrus; AG = angular gyrus.

There were no LH ROIs that demonstrated a characteristic response for NOV/MET that was significantly higher than the response for FAM/MET. Instead, brief periods of statistically higher HbO values for NOV/MET versus FAM/MET likely have other explanations, including the absence of a characteristic initial dip in HbO in LIFGtri (from 1-3s) and LSTG (from 1-5s), an atypical uptick in HbO values at the end of the time series for LIFGop (from 25-29s), and a nearly flat, noncanonical response in LSTG and LMTG for NOV/MET. In contrast, three RH ROIs were activated for NOV/MET but not FAM/MET: RIFGop, RMTG, and RAG. In part due to the large and unexpected initial decreases in HbO in these ROIs for NOV/MET, periods where values statistically differed from FAM/MET were brief, from 15-18s and 24-29s in RIFGop, 0-4s and 14-18s in RMTG, and 16-17s in RAG.

Several activated LH regions had significantly greater increases in HbO for FAM/MET than NOV/MET, including LMFG (from 2-30s, *q* < 0.05), LIFGtri briefly (from 17-18s, *q* < 0.05), LMTG (from 3-15s, *q* < 0.05), LITG (from 0-12s, *q* < 0.05), and LSMG (from 5-9s, *q* < 0.05). In terms of RH regions, increases in HbO were significantly higher for FAM/MET compared to NOV/MET in RMFG (from 6-20s and 28s, *q* < 0.05) and RIFGtri (from 7-13s and 17s, *q* < 0.05). The full t-test results comparing data from the NOV/MET and FAM/MET conditions are reported in Supplemental Table 4.

### 3.2. Aim #2: Relationships between brain activity and task behavior

#### 3.2.1. Statistical comparison of behavioral measures by condition

Compared to the random effects-only model, the model of interest was significantly better at predicting RT than the base random-effects-only model (χ² = 121.56, *p* < 0.001). Within the model of interest, RTs were significantly longer for NOV/LIT (β = 0.42, *SE* = 0.06, *t* = 6.641, *p* < 0.001), FAM/MET (β = 0.34, *SE* = 0.06, *t* = 5.334, *p* < 0.001), and NOV/MET (β = 0.92, *SE* = 0.06, *t* = 14.368, *p* < 0.001) compared to the reference condition (i.e., FAM/LIT). Post-hoc tests were significant for every comparison except for the comparison of NOV/LIT to FAM/MET (β = 0.08, *SE* = 0.06, *t* = 1.292, Tukey-adjusted *p* = 0.570). See Figure 7A for a visualization of model effects and Table 2 for the complete pairwise comparison findings.

**Figure 7.**
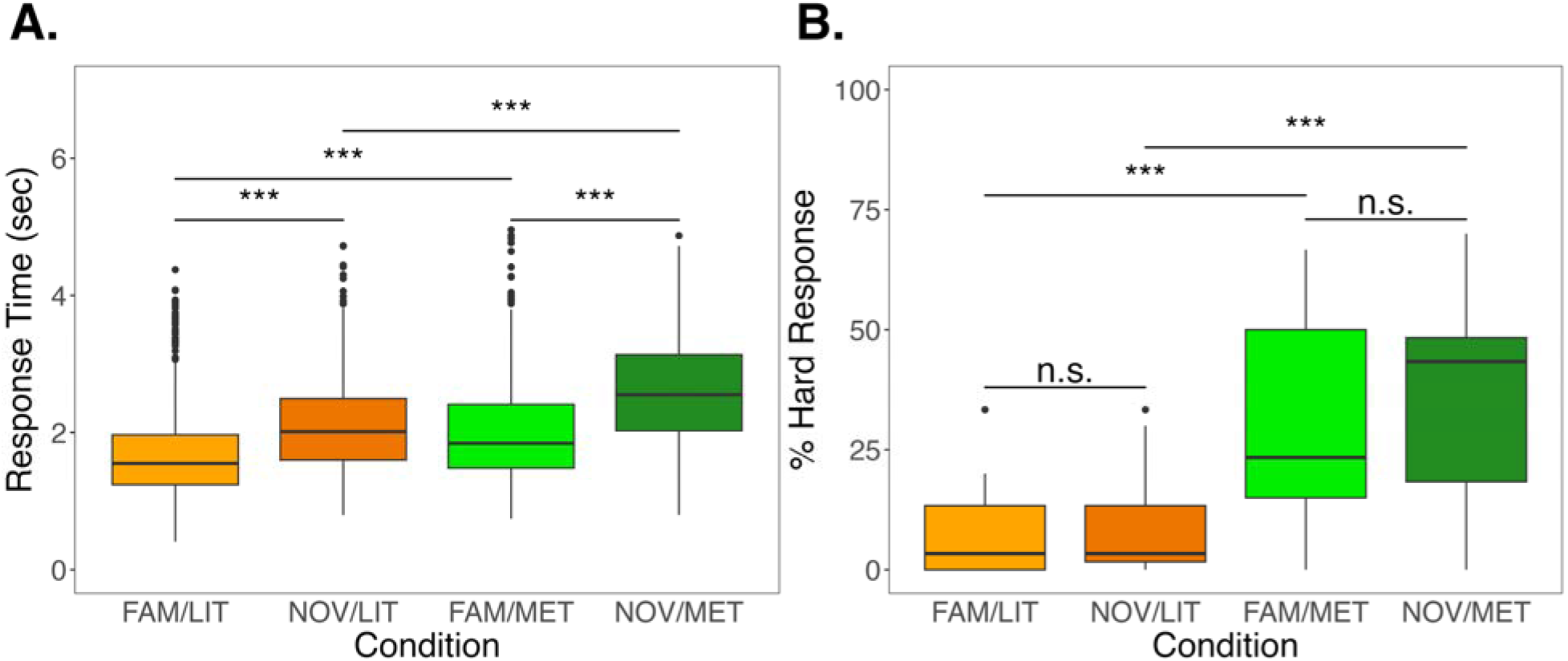
Between-condition differences in the A) button press reaction times (RTs) and B) percentage of “hard” key responses during the fNIRS task. Effects reflect Pairwise comparisons were evaluated using least squares means with Tukey *p*-value adjustment, \*\*\**q* < 0.001 or not significant (n.s.) Abbreviations: FAM/LIT = familiar literal sentences, NOV/LIT = novel literal sentences, FAM/MET = familiar metaphors, and NOV/MET = novel metaphors.

**Table 2.** Estimated Marginal Means from the Linear Mixed Effects Model Predicting Reaction Time by Condition.

| Comparison | Estimate | Standard Error | *t*-Statistic | *q*-value |
| --- | --- | --- | --- | --- |
| FAM/LIT vs. NOV/LIT | -0.424 | 0.065 | -6.55 | < 0.001*** |
| FAM/LIT vs. FAM/MET | -0.340 | 0.065 | -5.26 | < 0.001*** |
| FAM/LIT vs. NOV/MET | -0.917 | 0.065 | -14.18 | < 0.001*** |
| NOV/LIT vs. FAM/MET | 0.083 | 0.065 | 1.29 | 0.570 |
| NOV/LIT vs. NOV/MET | -0.493 | 0.065 | -7.63 | < 0.001*** |
| FAM/MET vs. NOV/MET | -0.577 | 0.065 | -8.92 | < 0.001*** |

Notes: Pairwise comparisons of the significant effect of condition on reaction times from the fNIRS experiment were evaluated using least squares means with Tukey *p*-value adjustment, \*\*\**q* < 0.001. Abbreviations: FAM = familiar, NOV = novel, LIT = literal, MET = metaphorical.

Similar to RT, the model of interest for key response had significantly higher predictive power than the random-effects-only model (χ² = 76.22, *p* < 0.001). Compared to FAM/LIT, there was a significantly higher number of “hard” key responses for FAM/MET (β = 2.48, *SE* = 0.37, *t* = 6.683, *p* < 0.001) and NOV/MET (β = 2.83, *SE* = 0.06, *t* = 7.627, *p* < 0.001) but not NOV/LIT (β = 0.51, *SE* = 0.39, *t* = 1.316, *p* = 0.188). Post-hoc tests revealed each comparison of metaphorical and literal sentences was significant (see Figure 7B and Table 3), but the number of “hard” key responses did not significantly differ between familiar and novel phrases for either literal (β = -0.51, *SE* = 0.39, *t* = -1.316, Tukey-adjusted *p* = 0.553) or metaphorical (β = -0.353, *SE* = 0.32, *t* = -1.110, Tukey-adjusted *p* = 0.684) sentences.

**Table 3.**
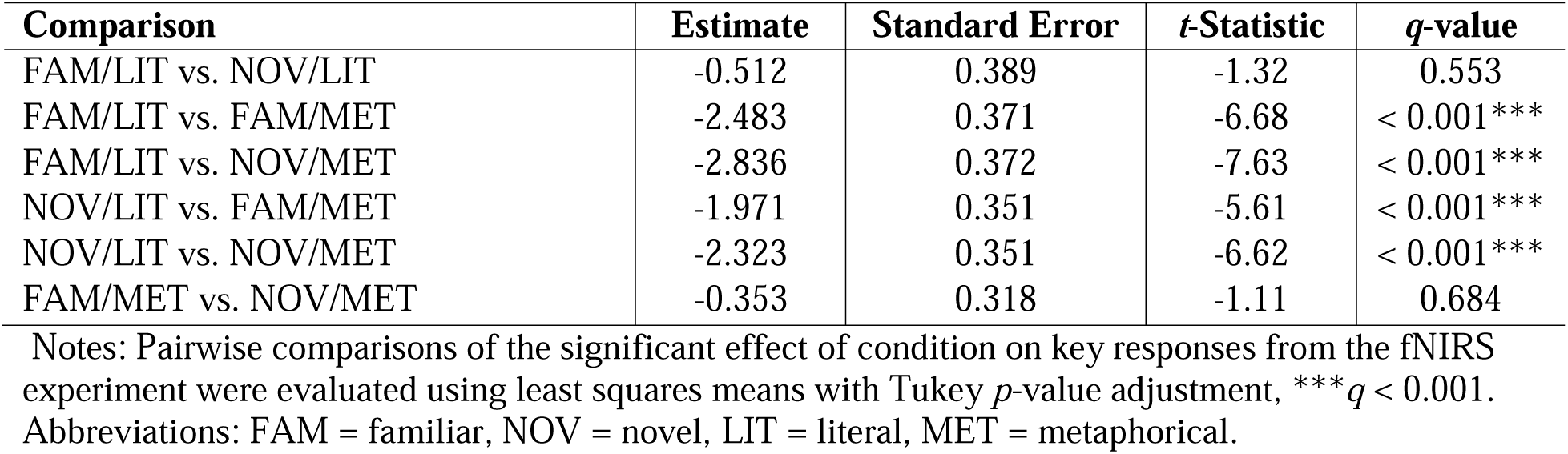
Estimated Marginal Means from the Logistic Mixed Effects Model Predicting Key Response by Condition.

#### 3.2.2. Relationships between activation within regions of interest and task performance

Due to a PsychoPy saving error, five participants had missing behavioral data from the fNIRS runs and were excluded from the LASSO regressions. For the remaining participants, the only variable retained in the LASSO regression predicting the difference in RT between MET and LIT conditions was LMFG (β = -0.485, 95% CI [-0.798, 3.422], *Z* = -2.274, *p* = 0.60). The LASSO regression predicting the difference in the percentage of “hard” key responses between the MET and LIT conditions resulted in a null model.

One variable was retained in the LASSO regression predicting the difference in RT between the NOV/MET and FAM/MET conditions, LMTG (β = 0.432, 95% CI [-0.524, 0.842], *Z* = 2.025, *p* = 0.174). Like the broader contrast, the LASSO predicting the difference in the percentage of “hard” key responses between NOV/MET and FAM/MET resulted in an empty model.

## 4. Discussion

This study was motivated by outstanding questions regarding the role of novelty in the recruitment of neural resources during metaphor processing and the extent to which neurologically healthy young adults rely on regions outside the canonical left lateralized language network for tasks that are hypothesized to be more difficult (e.g., novel over familiar phrases, metaphors versus literal sentences). Using fNIRS and a modified version of the paradigm employed by Cardillo et al. (2012), we found that activation was significantly greater for metaphorical than literal sentences across several frontal and parietal ROIs—with greater condition differences within the LH than RH—whereas greater changes in HbO were seen in superior and middle temporal cortex for literal sentences over metaphors. Contrary to expectations, we found that activity across several LH ROIs was significantly greater for familiar than novel metaphors, whereas the opposite (i.e., higher activation for NOV/MET > FAM/MET) was true of specific RH ROIs, albeit with generally weak effects. Participants judged metaphors to be significantly harder to understand than literal phrases, regardless of familiarity. Conversely, reaction times were condition-dependent, varying by both familiarity and sentence type and relating to between-condition differences in activation within certain ROIs. We discuss each finding in greater detail below and couch our results within the existing literature.

### 4.1. Greater activity for metaphors over literal sentences occurs within a widespread network of bilateral, but left dominant frontal and parietal regions

Theories regarding the neural correlates of figurative versus non-figurative language historically revolve around the laterality of activity for different types of stimuli. Importantly, all metaphors in this investigation could reasonably be classified as more novel than familiar—and certainly not conventionalized—given that even the “familiar” phrases (i.e., FAM/MET and FAM/LIT) were novel sentences that the participants were exposed to only once, just before completing the fNIRS experiment. Thus, our findings for the broad MET versus LIT comparison generally align with both coarse semantic coding theory (Beeman, 1998; Beeman et al., 1994; Jung-Beeman, 2005) and the graded salience hypothesis (Giora, 1997, 2003; Giora et al., 2000). Specifically, as expected, literal sentences were processed predominantly within the LH, and metaphors were also processed across perisylvian LH regions. Metaphors resulted in greater RH activation than literal sentences in three ROIs (RIFGtri for a brief early time window, RIFGop, and RAG) compared to greater RH activation in only one ROI (RMTG) for literal versus metaphorical sentences.

#### 4.1.1. Right hemisphere activity differences between metaphorical and literal sentences

Despite these general trends, it is important to acknowledge nuances in the findings, particularly for the RH findings. While increases in HbO within RIFGop and RAG were observed for the MET but not LIT condition, the effects had a small magnitude and relatively short duration (from ∼15-20s, see Figure 4), meaning that these regions were only weakly activated in the middle of the 30-second block across participants. Given that both RIFGop and RAG are implicated in domain-general cognitive control (Fedorenko et al., 2013), it may be that these regions were engaged in conjunction with their LH homologues only after participants realized their difficulty in processing metaphorical stimuli, a point we return to later in the discussion. The more notable RH finding was the recruitment of RIFGtri for both conditions, with a slightly larger response for MET over LIT. This finding is consistent with most prior meta-analyses of metaphor processing (Bohrn et al., 2012; Reyes-Aguilar et al., 2018; Yang, 2014) that reported activations in RIFG for the contrast of all metaphors (regardless of novelty/familiarity) to literal sentences. Importantly, however, RIFGtri was the only RH regions that demonstrated strong activation across participants, and the effects were not statistically different between MET and LIT conditions across most time windows. Thus, our comparisons of metaphorical and literal sentences do not lend strong support of RH dominance for metaphor processing.

#### 4.1.2. Left hemisphere activity for metaphors versus literal sentences: Domain-specific explanations

In contrast to the RH results, both metaphorical and literal sentences activated nearly all left frontal and temporal ROIs traditionally linked to language processing, including LIFGtri, LIFGop, LSTG, LMTG, and LITG. The starkest LH difference for metaphors versus literal sentences was observed in parietal regions: while only early and weak effects within LSMG and LAG were seen for literal sentences, large and sustained effects within these ROIs (as well as within LITG) were evident for metaphors. There is some consensus that left inferior parietal regions play a central role in the integration of sensory information (Binder & Desai, 2011; Lambon Ralph et al., 2017; Xu et al., 2017). Traditionally, LSMG has been implicated in phonological rather than semantic processes, a role supported by numerous lesion-symptom mapping studies in patients with aphasia and phonological impairments (Alyahya et al., 2020; Chen et al., 2019; Dell et al., 2013; Fridriksson et al., 2016, 2018; Halai et al., 2017; Henseler et al., 2014; Lacey et al., 2017; Mirman, Chen, et al., 2015; Mirman, Zhang, et al., 2015; Pillay et al., 2014; Schumacher et al., 2019; Schwartz et al., 2012; Tochadse et al., 2018) as well as meta-analytic evidence from functional imaging (Hodgson et al., 2021; Humphreys & Lambon Ralph, 2015; Vigneau et al., 2006). LSMG is activated when phonological working memory or decoding demands are high, as in the case of re-reading or reading difficult texts (Martin et al., 2015; Vigneau et al., 2006). In our study, in an effort to better understand the metaphors, participants may have read each phrase multiple times during the 5s trial window or perhaps even sub-vocally read the phrases. While neither explanation can be confirmed, the written nature of our stimuli and associated reading processes provide a logical account for the LSMG difference between conditions.

On the other hand, the classic language model (Geschwind, 1970) positions LAG as the seat of semantic concepts—particularly for visual words. More recent proposals suggest that the anterior temporal lobe (ATL) and LAG serve as dual semantic hubs or convergence zones, with LAG particularly tasked with processing thematic relations between concepts, performing combinatorial semantics, or perhaps heteromodal integrative processes (Binder et al., 2009; Binder & Desai, 2011; Bonner et al., 2013; Mirman et al., 2017; Price et al., 2015; Seghier, 2013). Based on meta-analytic results from 386 studies, (Humphreys & Lambon Ralph, 2015) concluded that task positive activations in LAG are associated with episodic retrieval and sentence-level processing while activations while more dorsal activity (in the intraparietal sulcus/ventral superior parietal lobule) occur when tasks require executive semantic decisions. Theories regarding the nature of new metaphor learning and comprehension (Bowdle & Gentner, 2005; Gentner & Bowdle, 2008; Jung-Beeman, 2005) indicate that understanding novel metaphors requires retrieval of context-specific semantic knowledge and fast mapping of abstract target to concrete source concepts. Thus, the likelihood that metaphors necessitate a higher semantic relational load than literal sentences may explain the LAG condition differences.

Differences between metaphors and literal sentences in the nature and degree of semantic load may also explain dissociations in temporal and LIFGtri activation patterns between conditions. More specifically, as shown in Figure 3, LpSTG, LpMTG, and LpITG were activated for metaphors while the anterior to mid LSTG and LMTG were not, whereas the opposite was true for literal sentences. LIFGtri was also activated in both conditions but to a significantly greater extent for metaphorical sentences. Evidence from multiple individual studies and meta-analyses (Davey et al., 2015, 2016; Hodgson et al., 2021; Jackson, 2021; Krieger-Redwood & Jefferies, 2014; Noonan et al., 2013) indicates that LIFGtri and LpMTG (with extension to LpITG) are specifically required for tasks that require high versus low degrees of semantic control, i.e., the ability to access and manipulate context-specific conceptual information. Multiple models of semantic cognition feature the anterior temporal lobe (including the temporal pole, extending to anterior middle and inferior temporal gyri) as either a graded amodal (Humphreys & Lambon Ralph, 2015; Lambon Ralph et al., 2017; Xu et al., 2017) or taxonomic (Mirman et al., 2017) semantic hub, with a leftward bias for linguistic information and a rightward bias for nonverbal, visual, and social tasks (Rice et al., 2015). Extrapolating to the current study, it is reasonable to assume that the bilateral (but LH dominant) MTG activation for literal sentences reflects more automatic retrieval of lexical-semantic information whereas greater LIFGtri and left posterior temporal activation for metaphors reflects higher semantic control demands for this sentence type.

Our left frontal and temporal findings are largely consistent with prior fMRI studies of metaphor as all prior meta-analyses (Bohrn et al., 2012; Huang et al., 2023; Rapp et al., 2012; Reyes-Aguilar et al., 2018; Yang, 2014) reported activation within parts of LIFG—particularly LIFGtri—and LpSTG and LpMTG extending to LpITG for metaphors over literal stimuli. However, relatively few parietal findings were reported in past meta-analyses of metaphorical and figurative language processing, in contrast with our results. While Rapp et al. (2012) and Yang (2014) both reported LSMG in their contrast of all metaphorical over literal stimuli, LAG was found only by Yang (2014), and other reviews (Bohrn et al., 2012; Huang et al., 2023; Reyes-Aguilar et al., 2018) found no inferior parietal activations for this comparison. Given the nature of our stimuli, the contrast of novel metaphors over literal stimuli is a closer match to our comparisons, yet of the meta-analyses that compared similar conditions (including Bohrn et al., 2012; Huang et al., 2023; Rapp et al., 2012), only Yang (2014) reported left parietal activity (within LSMG) for this contrast. Furthermore, in the study on which ours was based, Cardillo et al. (2012), the authors did find inferior parietal activation for metaphors, but they reported highly right lateralized, not LH-dominant parietal activity.

#### 4.1.3. Left hemisphere activity for metaphors versus literal sentences: Domain-general explanations

An alternative, domain-general hypothesis to our above domain-specific (phonological and semantic) explanations for LSMG and LAG activation may better resolve discrepancies in past and present parietal findings. Through the use of functional localizers across several studies, Fedorenko and colleagues (Duncan, 2010; Fedorenko et al., 2011, 2012, 2013) distinguished language-specific frontotemporal regions that comprise a core, domain-specific network from the bilateral domain-general, multiple demand (MD) network, which encompasses many regions we found activated for metaphors but not literal sentences, including the posterior inferior frontal sulcus, IFGop, and the dorsal inferior parietal cortex. Although several studies have demonstrated that the MD network is not highly engaged in language processing when task demands are kept low (Blank & Fedorenko, 2017; Diachek et al., 2020; Fedorenko et al., 2011; Wehbe et al., 2021), there is some evidence that MD and language network regions jointly come online under specific conditions, such as for linguistic manipulations (Diachek et al., 2020; Kuperberg et al., 2003; Mollica et al., 2020; Rodd et al., 2005). In a large-scale study including 481 participants, (Diachek et al., 2020) reported that bilateral MD network ROIs strongly responded to explicit language tasks—particularly visual tasks involving sentence ratings or plausibility judgments—but not passive listening or reading paradigms, whereas language network ROIs responded equally to explicit and passive tasks. Our study required participants to judge the ease with which they understood presented sentences during the fNIRS task. As such, these findings may partially explain why we found activation within LSMG and LAG as well as RIFGop and RAG for metaphors, but they cannot account for the lack of such activity for literal sentences, given that we required participants to complete the comprehensibility judgment on all sentences in every condition.

It is possible that the twin demands of the explicit judgment and higher processing difficulty resulted in higher MD network engagement for metaphors over literal sentences, at least for some individuals. In this vein, while LMFG was not significantly activated at a group level for metaphors (or literal sentences), participants who exhibited more similar RTs between the two conditions activated LMFG to a greater extent for the MET than LIT condition than participants who experienced a greater RT condition difference. In other words, participants who showed this effect may have recruited LMFG to exert top-down executive control over other regions or to perform other cognitive processes (e.g., working memory, attention) that resulted in their processing speed for metaphorical sentences to closely resemble that of literal sentences. While plausible, this explanation cannot be confirmed, especially given that activations within LMFG for metaphorical sentences were spatially localized to both dorsal LMFG—which is traditionally included as part of the “core” language network—as well as more rostral LMFG, which is part of the MD network.

### 4.2. Greater activity is seen for (more) familiar, rather than completely novel, metaphors

Consistent with the broad contrast findings, several regions were significantly recruited for processing both novel and (more) familiar metaphors, including bilateral IFGtri (although LH > RH for NOV/MET), LIFGop, LITG, LSMG, and LAG. Contrary to expectations, very few regions were significantly activated more in the NOV/MET than FAM/MET condition, and all effects in this direction either had an uncharacteristic shape to the hemodynamic response (i.e., LSTG and RIFGop) or were of a short duration and magnitude (i.e., bilateral IFGop, RMTG, RAG). That said, most responses that likely represented a true HRF difference between conditions were contained in the RH. Interestingly, these ROIs were ones that we hypothesized were recruited for the MET > LIT contrast to assist their LH homologues in retrieval of lexical-semantic representations (RMTG) or relational semantics, semantic control, or domain-general executive control (RAG and RIFGop), arguably all of which would be needed to process novel metaphors for the first time. More generally, these findings provide some support—albeit weak—for the hemispheric laterality difference for novel over (more) familiar metaphors proposed by the coarse semantic coding theory (Beeman, 1998; Beeman et al., 1994; Jung-Beeman, 2005) and the graded salience hypothesis (Giora, 1997, 2003; Giora et al., 2000). They also generally agree with the conclusion drawn by (Yang, 2014) in their investigation of task paradigm influences on activation patterns: they concluded that the RH is only involved when the metaphor was novel, when the meaning was presented in a sentential context, and when the task was a semantic relatedness judgment.

In addition to the ROIs listed above, significant increases in HbO were seen in bilateral MFG and LMTG for the FAM/MET condition, and activations were greater for FAM/MET versus NOV/MET in these regions—as well as within RIFGtri and trending for LIFGtri. Given the uniqueness of the design by Cardillo et al. (2012) that we modified for our study, no contrast investigated in prior neuroimaging meta-analyses on metaphor is similar to the FAM/MET > NOV/MET comparisons here. Intriguingly, however, the activation patterns we observed for our comparison are most similar to patterns reported for the contrast of novel over conventional metaphors, which included bilateral/left MFG (Bohrn et al., 2012; Huang et al., 2023; Rapp et al., 2012), RIFGtri/RIFGorb (Huang et al., 2023; Rapp et al., 2012), LIFGtri/LIFGop (Bohrn et al., 2012; Rapp et al., 2012), and LpMTG/LpITG (Bohrn et al., 2012). Major differences between these findings and ours were activations reported in meta-analyses that are outside our montage (e.g., left parahippocampal gyrus, RdACC, right insula, LmPFC) but also activity within bilateral or right SFG observed in certain prior studies (Huang et al., 2023; Rapp et al., 2012). Surprisingly, our results are also the inverse of those reported by Cardillo and colleagues (2012). Applying a parametric modulation of familiarity, Cardillo et al. (2012) found that activation within bilateral IFGtri, LpMTG, and right posterolateral occipital gyrus *decreased* as familiarity increased, whereas our results suggest that activation within these regions *increases* with increased familiarity.

Initially, these collective findings seem at odds but could be reconciled when the likely computations of the implicated regions are considered in conjunction with the differences between our task paradigm and that of Cardillo et al. (2012). The Career of Metaphor (Bowdle & Gentner, 2005) posits that the number of encounters influences the manner in which a novel metaphor is processed, with increasing exposures resulting in eventual semantic categorization of the metaphor as a learned single unit. Until that point, the learning and mapping process would arguably place high demands on semantic control and/or domain-general executive resources, consistent with activation in LIFGtri, LpMTG, and bilateral MFG in particular (Fedorenko et al., 2011, 2013; Jefferies, 2013; Lambon Ralph et al., 2017). In Cardillo et al. (2012), participants were pre-exposed to two-thirds of the metaphors prior to the fMRI experiment, with 40 of 120 metaphors presented five times in advance and 40 of 120 presented twice. In our study, the list of stimuli included metaphors and literal sentences, and participants were pre-exposed to half of each sentence type (i.e., 30 metaphors and 30 metaphors) only three times each before the fNIRS task. It may be that a certain number of exposures of completely novel—and frankly, unusual— metaphors are needed for participants to successfully map and truly process such phrases. If that is the case, then our overall weaker—but spatially similar—effects for NOV/MET compared to FAM/MET (see Figure 5) make sense. Moreover, the fewer pre-exposures for a smaller percentage of stimuli in our study compared to Cardillo et al. (2012)’s may have resulted in the FAM/MET stimuli not reaching a critical learning mass, thereby requiring a higher degree of semantic and cognitive control resources than expected.

This explanation gains traction when the relationships between task behavior and activation patterns for NOV/MET and FAM/MET are considered. Specifically, we found that a greater difference between the NOV/MET and FAM/MET RTs was associated with greater positive changes in HbO for LMTG, likely LpMTG based on image reconstructions (Figure 5). Slower RTs for NOV/MET compared to FAM/MET may reflect attempts at truly comprehending the completely novel metaphors. If true, it is likely that participants with longer RTs would rely on LpMTG for controlled retrieval of the correct word meanings necessary to successfully map the source to the target concepts for metaphorical phrases that they needed to process quickly (within 5s) for which they had no previous exposure. While this interpretation cannot be confirmed by our methods, a concrete conclusion that can be drawn is that difference in novel versus (more) familiar processing speed was linked to activation not within the RH (as many metaphor theories would suggest) nor in traditional domain-general MD regions. Overall, this work highlights the importance of continuing to resolve questions about the recruitment of resources in the processing of harder linguistic stimuli outside of the language-specific brain especially in processing high-difficulty semantic content, such as metaphors.

### 4.3. Study limitations

There are several important limitations to the current study due to technical constraints of fNIRS. First, the measurement depth of fNIRS is approximately 1.5 cm into the cortex. Thus, activation in deep gray matter structures cannot be acquired using fNIRS, which prevented us from examining possible contributions of regions found in prior studies of metaphor (Bohrn et al., 2012; Yang, 2014) such as the mPFC, insula, and dACC. Relatedly, our montage did not span the entire cortical surface, meaning that we did not have coverage of certain superficial areas implicated in previous studies (e.g., Cardillo et al., 2012), such as the anterior IFGorb and posterolateral occipital gyrus. The continuous wave fNIRS system used in this study was not a high-density optical tomography array, so spatial resolution was limited by our sparse probe. Furthermore, while our procedures involved careful head measurements and placement of the cap relative to fiducials and participants’ external anatomy, we did not digitize optode locations. Our method of assigning ROI labels to channels has been used previously in published studies (Gilmore et al., 2021; Li et al., 2020), but groupings were made according to approximate, likely anatomical regions. Because of these collective impacts on spatial precision, we did not attempt a more fine-grained parcellation of ROIs beyond the AAL labels, yet doing so meant that we aggregated data across large swaths of tissue. This approach led to a blurring of effects in certain regions, which was particularly evident in activation differences observed for anterior to mid versus posterior temporal regions for conditions of interest (see Figures 3 and 5).

While many researchers approach language research by excluding participants who are not English monolinguals, 60% of the world speaks more than one language (Grosjean, 2024; Haspelmath, 2005), and a bilingual is not the sum of two monolinguals (Grosjean, 1989). The inclusion of bi/multilingual individuals adds complexity to the interpretation of our data, but it is also important to study language in a manner that can be generalized to the greater population. It is possible that another sample with fewer bi/multilingual participants may have yielded different results, particularly with regards to the novel versus familiar metaphor findings as vocabulary size can impact semantic processing. However, while we did not obtain external measures that can reliability index language proficiency, all participants indicated that English was a primary language. One potential future direction would be to collect and use more specific language proficiency measures as covariates when analyzing fNIRS data in a bi/multilingual sample.

There were also some study limitations with regards to metaphor theory. Within conceptual mapping theory (Lakoff, 2014; Lakoff & Johnson, 1980), there are three main metaphorical types in English, yet our stimuli were limited to conceptual/structural metaphors. Orientational (e.g., HAPPY IS UP, SAD IS DOWN) or ontological (e.g., IDEAS ARE OBJECTS, INFLATION IS AN ENTITY) metaphors may generate different activation patterns. Moreover, to move beyond the single most studied sentence structure in metaphor studies (i.e., nominal phrases, (Holyoak & Stamenković, 2018), our stimuli included half nominal and half predicate phrases as well as base terms that were either auditory or motion-based. While this approach may have accounted for several variables highlighted by (Koller et al., 2022) as uncontrolled influences in past studies, it also means that our results may differ from many prior investigations for various reasons.

### 4.4. Future directions

An important future direction for this work is moving beyond young, neurologically healthy adult samples. Age likely contributes to different activation patterns for metaphor and figurative language processing due to differences in brain function, conceptual knowledge, and executive skills between older and younger adults (Cabeza, 2001; Cabeza et al., 2002; N. A. Dennis & Cabeza, 2011; Reuter-Lorenz & Cappell, 2008), yet most metaphor neuroimaging studies are limited to younger individuals. Metaphors and other figurative language processing abilities (including irony and sarcasm) have been characterized as impaired across a variety of clinical disorders, including Alzheimer’s disease (Rapp & Wild, 2011), schizophrenia (Sponheim, 2003), autism (Wang et al., 2006), both right and left hemisphere stroke (Tompkins, 1990), and traumatic brain injury (M. Dennis & Barnes, 1990). Simultaneously studying individuals with different clinical conditions could highlight shared and distinct behavioral and neural factors that lead to such difficulty, thereby further clarifying the neurobiology of metaphor.

## 5. Conclusions

Metaphors are a profound way of understanding, making sense of, and discussing novel experiences, making them a vital part of human sociocultural functioning and sociocultural evolution (Lakoff, 2014). Yet, metaphors are still poorly understood, especially with respect to their neural underpinnings. Understanding what additional neural resources are associated with novel metaphor processing provides important insights into leveraging existing semantic representations to make sense of novel contexts for which conventionalized descriptions do not yet exist. This study sought to deepen our understanding of how novel metaphors are processed in the brain by replicating and extending theoretically driven neuroimaging research done with fMRI (Cardillo et al., 2012). While our findings lend some credence to theories (Giora, 1997; Jung-Beeman, 2005) suggesting that metaphors—particularly novel metaphors—do engage RH cortex more than other phrase types (literal sentences, more familiar metaphors), our RH effects were less robust than condition differences within canonical left language network ROIs and domain-general MD regions. Activation patterns for the comparison of completely novel metaphors to more familiar metaphors (to which participants were exposed just prior to the fNIRS task) suggest a core set of regions (including bilateral IFGtri and LpMTG) comes online during semantic mapping of such phrases. When considering the present study results in light of past findings (particularly from Cardillo et al., 2012), it may be individuals must have a certain degree of exposure to novel metaphors before successful mapping of these metaphors is substantiated in the brain.

## Glossary

activation likelihood estimation: ALE
angular gyrus: AG
anterior cingulate cortex: ACC
anterior temporal lobe: ATL
Automated Anatomical Labeling: AAL
Brodmann area: BA
deoxyhemoglobin: HbR
dorsal: d
false discovery rate: FDR
familiar: FAM
functional near-infrared spectroscopy: fNIRS
general linear model: GLM
inferior frontal gyrus, pars opercularis: IFGop
inferior frontal gyrus, pars orbitalis: IFGorb
inferior frontal gyrus, pars triangularis: IFGtri
inferior temporal gyrus: ITG
Least Absolute Shrinkage and Selection Operator: LASSO
left: L
left hemisphere: LH
literal: LIT
medial: m
metaphorical: MET
middle frontal gyrus: MFG
middle temporal: MTG
Montreal Neurological Institute: MNI
multiple demand: MD
novel: NOV
optical density: OD
oxyhemoglobin: HbO
posterior: p
prefrontal cortex: PFC
reaction time: RT
regions of interest: ROIs
right: R
right hemisphere: RH
supramarginal gyrus: SMG
superior frontal gyrus: SFG
superior temporal: STG

## Acknowledgments

We extend our thanks to other members of The Aphasia Network (TAN) Lab who assisted with participant recruitment, testing, and fNIRS data collection sessions. We are also grateful to our participants for their time spent in completing study procedures.

## Author contributions: CRediT

**Anna Schwartz:** Conceptualization, Data curation, Formal analysis, Investigation, Methodology, Visualization, Writing – original draft; **Natalie Gilmore:** Investigation, Methodology, Writing – review and editing; **Erin Meier:** Conceptualization, Funding acquisition, Data curation, Formal analysis, Investigation, Methodology, Project administration, Supervision, Visualization, Writing – original draft

## Declaration of Generative AI in Scientific Writing and Figures

During the preparation of this work the senior author (E.M.) used ChatGPT-5 to identify and correct errors in the R code that was used to specify statistical models and generate the figures. This approach was used in lieu of using other online resources. After using this tool, the senior author reviewed and edited the suggested code as needed; the authors take full responsibility for the content of the published article. No generative AI or AI-assisted technologies were used to generate any text within the manuscript or to manipulate or edit any figures.

## Footnotes

1 During intake, participants were asked to provide their “gender/sex” in a blank textbox. Because the interpretation of this prompt and their response was left to the discretion of the participants, we cannot assume this variable corresponds to biological sex for all individuals. Thus, this variable was balanced across participants but not investigated further in statistical analyses.

2 The number of datapoints within each 1s window is based on the sampling rate at which fNIRS data were collected (i.e., 11.2 Hz).

