## Supplemental Tables 1-4 for "The neural correlates of novel versus familiar metaphors in healthy young adults: A functional near-infrared spectroscopy study"

Anna Schwartz, PhD, CF, MS-SLP^1,2^, Natalie M. Gilmore, PhD, CCC-SLP^3,4^, Erin L. Meier, PhD, CCC-SLP^1^

^1^Department of Communication Sciences and Disorders, Northeastern University, Boston, MA, USA

^2^Department of Physical Therapy, Movement, and Rehabilitation Sciences, Northeastern University, Boston, MA, USA

^3^Research Service, James A. Haley Veterans Hospital, Tampa, FL, USA

^4^Department of Neurosurgery, Brain, and Spine, Morsani College of Medicine, University of South Florida, Tampa, FL, USA

**Supplemental Table 1. Demographic and Behavioral Data for All Participants (*n* = 28)**

| **ID** | **Gender** | **Age**  **(years)** | **Education**  **(years)** | **Handedness** | **Race** | **Ethnicity** | **Native**  **Language** | **Language**  **Status** | **MMSE**  **(/30)** | **NIHTB**  **PVT**  **(Age-Adj**  **SS)** | **NIHTB**  **PCPT**  **(Age-Adj**  **SS)** | **NIHTB**  **FICAT**  **(Age-Adj**  **SS)** | **NIHTB**  **DCCS**  **(Age-Adj**  **SS)** | **Base**  **Term**  **Yeses**  **(%)** |
| --- | --- | --- | --- | --- | --- | --- | --- | --- | --- | --- | --- | --- | --- | --- |
| P1 | M | 23 | 16 | R | White | Non-Hispanic | English | Bilingual | 30 | 123 | 103 | 98 | 120 | 100.00 |
| P2 | W | 26 | 16 | R | White | Non-Hispanic | English | Multilingual | 29 | 116 | 127 | 89 | 93 | 100.00 |
| P3 | W | 18 | 13 | Ambi-  dextrous | Middle  Eastern | Persian/Iranian | English | Multilingual | 29 | 116 | 111 | 90 | 105 | 100.00 |
| P4 | W | 21 | 15 | R | Biracial  (White/  Asian) | Non-Hispanic | English | Monolingual | 30 | 136 | 102 | 108 | 107 | 98.00 |
| P5 | W | 18 | 15 | R | Indian/  Asian | Indian/  Non-Hispanic | English | Bilingual | 29 | 116 | 122 | 101 | 117 | 100.00 |
| P6 | W | 19 | 14 | R | White | Non-Hispanic | Russian | Bilingual | 29 | 115 | 128 | 102 | 117 | 100.00 |
| P7 | W | 23 | 16 | R | White | Caucasian | N/A | N/A | 30 | 129 | 92 | 104 | 114 | 100.00 |
| P8 | M | 25 | 17 | R | White | White | N/A | N/A | 29 | 128 | 54 | 105 | 109 | 100.00 |
| P9 | M | 29 | 19 | R | White | Non-Hispanic | English | Monolingual | 30 | 114 | 78 | 90 | 94 | 98.11 |
| P10 | W | 26 | 18 | R | Asian | Non-Hispanic | Hindi | Multilingual | 28 | 93 | 122 | 94 | 104 | 98.18 |
| P11 | W | 24 | 18 | R | Asian | Non-Hispanic | English | Bilingual | 28 | 60 | 137 | 99 | 103 | 88.46 |
| P12 | M | 23 | 18 | R | Asian | Non-Hispanic | English | Multilingual | 28 | 95 | 131 | 104 | 114 | 85.19 |
| P13 | M | 21 | 16 | R | White | Non-Hispanic | English | Bilingual | 29 | 125 | 91 | 97 | 130 | 98.18 |
| P14 | M | 26 | 17 | R | Asian | Non-Hispanic | Tamil | Bilingual | 30 | 110 | 76 | 78 | 64 | 76.92 |
| P15 | M | 25 | 22 | R | Asian | Non-Hispanic | English,  Marathi,  Hindi | Multilingual | 28 | 94 | 110 | 132 | 86 | 100.00 |
| P16 | W | 22 | 17 | L | Asian | Non-Hispanic | English | Multilingual | 29 | 95 | 119 | 92 | 108 | 98.18 |
| P17 | M | 23 | 16 | N/A | Asian | Non-Hispanic | Tamil,  English | Multilingual | 27 | 89 | 69 | 87 | 108 | 72.22 |
| P18 | W | 22 | 15 | R | Asian | N/A | English,  Marathi,  Hindi,  Gujrati | Multilingual | 29 | 90 | 80 | 76 | 73 | 78.18 |
| P19 | W | 23 | 17 | R | Asian | Non-Hispanic | English | Multilingual | 30 | 123 | 82 | 69 | 97 | 100.00 |
| P20 | W | 23 | 21 | R | Asian | Non-Hispanic | English | Multilingual | 30 | 100 | 120 | 87 | 85 | 100.00 |
| P21 | W | 25 | 20 | R | Asian | Non-Hispanic | English | Multilingual | 30 | 116 | 98 | 88 | 75 | 98.18 |
| P22 | M | 20 | 15 | R | White | Non-Hispanic | English | Multilingual | 30 | 132 | 90 | 108 | 129 | 100.00 |
| P23 | M | 28 | 22 | R | Asian | Non-Hispanic | English | Multilingual | 29 | 120 | 100 | 90 | 100 | 96.30 |
| P24 | M | 19 | 14 | R | White | Non-Hispanic | English | Multilingual | 29 | 109 | 123 | 102 | 123 | 100.00 |
| P25 | M | 23 | 18 | R | White | Non-Hispanic | English | Monolingual | 30 | 123 | 81 | 98 | 85 | 100.00 |
| P26 | W | 19 | 14 | L | White | Non-Hispanic | English | Monolingual | 29 | 98 | 128 | 96 | 123 | 94.55 |
| P27 | M | 29 | 18 | R | Asian | Non-Hispanic | English | Bilingual | 28 | 114 | 72 | 135 | 135 | 98.18 |
| P28 | M | 24 | 18 | R | Asian | Non-Hispanic | English | Multilingual | 28 | 83 | 75 | 71 | 69 | 88.89 |
| **AVG** |  | **23.11** | **16.96** |  |  |  |  |  | **29.07** | **109.36** | **100.75** | **96.07** | **103.11** | **95.28** |
| **STD** |  | **3.07** | **2.35** |  |  |  |  |  | **0.86** | **17.40** | **22.76** | **14.79** | **19.03** | **7.93** |

Notes: The National Institutes of Health Toolbox Tasks (NIHTB) each output an age-adjusted standard score (age-adj SS) that is calculated by comparing combined accuracy and reaction time information of the participant against normative samples^1–3^; scores from 85-115 are within normal limits. The percentage of “yes” responses on the base term task reflect the percentage of terms understood by a given participant. Abbreviations: M = man; W = woman; R = right; L = left; MMSE = Mini Mental State Examination^4^; PVT = NIHTB Picture Vocabulary Test; PCPS = NIHTB Pattern Comparison Processing Speed Test; FICAT = NIHTB Flanker Inhibitory Control and Attention Test; DCCS = NIHTB Dimensional Change Card Sort Test.

**Supplemental Table 2. All Experimental Stimuli by Phrase Type and Condition**

| **No.** | **Phrase Type** | **Nominal/ Predicate** | **Auditory/ Motion** | **Novelty** | **Base** | **Phrase** |
| --- | --- | --- | --- | --- | --- | --- |
| 1 | Literal | Predicate | Auditory | FAM | cheer | The fans cheered for the team. |
| 2 | Metaphor | Predicate | Auditory | FAM | cheer | The posters cheered for the candidate. |
| 3 | Literal | Predicate | Auditory | FAM | clash | His car's gears clashed in the cold. |
| 4 | Metaphor | Predicate | Auditory | FAM | clash | His smile clashed with his eyes. |
| 5 | Literal | Predicate | Auditory | FAM | coo | The gentle mom cooed at the baby. |
| 6 | Metaphor | Predicate | Auditory | FAM | coo | The cheap records cooed to the teenager. |
| 7 | Literal | Predicate | Motion | FAM | crawl | The spider crawled along the edge. |
| 8 | Metaphor | Predicate | Motion | FAM | crawl | The banker crawled through the contract. |
| 9 | Literal | Nominal | Motion | FAM | dig | The expedition was a desert dig. |
| 10 | Metaphor | Nominal | Motion | FAM | dig | The therapy was an archeological dig. |
| 11 | Literal | Nominal | Motion | FAM | dodge | His move was a quick dodge. |
| 12 | Metaphor | Nominal | Motion | FAM | dodge | His smile was a charming dodge. |
| 13 | Literal | Nominal | Auditory | FAM | flush | The only noise was a flush. |
| 14 | Metaphor | Nominal | Auditory | FAM | flush | His memoirs were a toilet flush. |
| 15 | Literal | Predicate | Auditory | FAM | growl | The panther growled at the photographer. |
| 16 | Metaphor | Predicate | Auditory | FAM | growl | The cruise ship growled at the fishing boat. |
| 17 | Literal | Nominal | Auditory | FAM | grunt | Her only input was a grunt. |
| 18 | Metaphor | Nominal | Auditory | FAM | grunt | The employee was a grunt. |
| 19 | Literal | Nominal | Auditory | FAM | hiss | The cat's reproach was a hiss. |
| 20 | Metaphor | Nominal | Auditory | FAM | hiss | His posture was a cat’s hiss. |
| 21 | Literal | Predicate | Auditory | FAM | howl | His lonely dog howled through the long night. |
| 22 | Metaphor | Predicate | Auditory | FAM | howl | His sore legs howled through the last mile. |
| 23 | Literal | Nominal | Motion | FAM | limp | The gait was a mild limp. |
| 24 | Metaphor | Nominal | Motion | FAM | limp | The winter was a heartbroken limp. |
| 25 | Literal | Predicate | Motion | FAM | lurch | The yellow taxi lurched through the lanes. |
| 26 | Metaphor | Predicate | Motion | FAM | lurch | The football team lurched through the season. |
| 27 | Literal | Nominal | Auditory | FAM | mumble | His advice was a mumble. |
| 28 | Metaphor | Nominal | Auditory | FAM | mumble | His handshake was a mumble. |
| 29 | Literal | Nominal | Motion | FAM | pounce | The cat's attack was a pounce. |
| 30 | Metaphor | Nominal | Motion | FAM | pounce | The purchase was a tiger pounce. |
| 31 | Literal | Predicate | Motion | FAM | puff | The housewife puffed up the down pillows. |
| 32 | Metaphor | Predicate | Motion | FAM | puff | The coach puffed up the football team. |
| 33 | Literal | Nominal | Motion | FAM | pull | The magnet was a weak pull. |
| 34 | Metaphor | Nominal | Motion | FAM | pull | The road was an irresistible pull. |
| 35 | Literal | Nominal | Auditory | FAM | purr | The kitten's reply was a purr. |
| 36 | Metaphor | Nominal | Auditory | FAM | purr | Her smile was a cat’s purr. |
| 37 | Literal | Predicate | Motion | FAM | retreat | The army retreated from the enemy lines. |
| 38 | Metaphor | Predicate | Motion | FAM | retreat | The painter retreated from his gloomy marriage. |
| 39 | Literal | Predicate | Auditory | FAM | sigh | The tailor sighed over the brass buttons. |
| 40 | Metaphor | Predicate | Auditory | FAM | sigh | The editorial sighed over the riots. |
| 41 | Literal | Predicate | Motion | FAM | slide | The sleigh slid down the hill. |
| 42 | Metaphor | Predicate | Motion | FAM | slide | The conversation slid into a wall. |
| 43 | Literal | Predicate | Motion | FAM | snake | The python snaked around its victim. |
| 44 | Metaphor | Predicate | Motion | FAM | snake | The lies snaked through her story. |
| 45 | Literal | Nominal | Auditory | FAM | squeal | The pig's protest was a squeal. |
| 46 | Metaphor | Nominal | Auditory | FAM | squeal | The bill was a corrupt squeal. |
| 47 | Literal | Predicate | Auditory | FAM | stutter | Her bashful suitor stuttered during dinner. |
| 48 | Metaphor | Predicate | Auditory | FAM | stutter | His feet stuttered on the dance floor. |
| 49 | Literal | Nominal | Motion | FAM | sweep | The chore was a quick sweep. |
| 50 | Metaphor | Nominal | Motion | FAM | sweep | The eviction was a mean sweep. |
| 51 | Literal | Nominal | Motion | FAM | tug | The puppy's grasp was a firm tug. |
| 52 | Metaphor | Nominal | Motion | FAM | tug | The shop display was a gentle tug. |
| 53 | Literal | Nominal | Motion | FAM | wave | The tsunami was a giant wave. |
| 54 | Metaphor | Nominal | Motion | FAM | wave | The letter was a goodbye wave. |
| 55 | Literal | Predicate | Auditory | FAM | whimper | The puppy whimpered in his pen. |
| 56 | Metaphor | Predicate | Auditory | FAM | whimper | The plants whimpered in the shadows. |
| 57 | Literal | Nominal | Auditory | FAM | whisper | The conversation was a hushed whisper. |
| 58 | Metaphor | Nominal | Auditory | FAM | whisper | His glance was a furtive whisper. |
| 59 | Literal | Nominal | Auditory | FAM | whistle | The referee's call was a whistle. |
| 60 | Metaphor | Nominal | Auditory | FAM | whistle | His smirk was a shameless whistle. |
| 61 | Literal | Predicate | Motion | NOV | balloon | The mattress ballooned to its full size. |
| 62 | Metaphor | Predicate | Motion | NOV | balloon | The kid’s courage ballooned during the fight. |
| 63 | Literal | Nominal | Auditory | NOV | buzz | The mosquito was an irritating buzz. |
| 64 | Metaphor | Nominal | Auditory | NOV | buzz | The ideas were a brain buzz. |
| 65 | Literal | Nominal | Motion | NOV | canter | The horse's trot was a canter. |
| 66 | Metaphor | Nominal | Motion | NOV | canter | His youth was a happy canter. |
| 67 | Literal | Predicate | Auditory | NOV | chant | The monks chanted in the mountains. |
| 68 | Metaphor | Predicate | Auditory | NOV | chant | The waves chanted to the surfer. |
| 69 | Literal | Nominal | Motion | NOV | clamber | The final ascent was an exhausting clamber. |
| 70 | Metaphor | Nominal | Motion | NOV | clamber | His work experience was a clumsy clamber. |
| 71 | Literal | Nominal | Auditory | NOV | crackle | The recording was a faint crackle. |
| 72 | Metaphor | Nominal | Auditory | NOV | crackle | The seizure was a brain crackle. |
| 73 | Literal | Predicate | Auditory | NOV | drum | The musician drummed in the park. |
| 74 | Metaphor | Predicate | Auditory | NOV | drum | The liquor drummed through his body. |
| 75 | Literal | Predicate | Motion | NOV | flounder | The giant seal floundered on the beach. |
| 76 | Metaphor | Predicate | Motion | NOV | flounder | The television show floundered in the Spring. |
| 77 | Literal | Nominal | Auditory | NOV | giggle | The child's answer was a giggle. |
| 78 | Metaphor | Nominal | Auditory | NOV | giggle | His ugly car is a giggle. |
| 79 | Literal | Nominal | Motion | NOV | glide | The skater's entrance was a glide. |
| 80 | Metaphor | Nominal | Motion | NOV | glide | The art major was a glide. |
| 81 | Literal | Predicate | Auditory | NOV | grunt | The boar grunted behind the orange trees. |
| 82 | Metaphor | Predicate | Auditory | NOV | grunt | The truck grunted at the small parking space. |
| 83 | Literal | Predicate | Auditory | NOV | hiss | The mean goose hissed at the gardener. |
| 84 | Metaphor | Predicate | Auditory | NOV | hiss | The designer purse hissed at the fakes. |
| 85 | Literal | Nominal | Motion | NOV | jog | The race course was an easy jog. |
| 86 | Metaphor | Nominal | Motion | NOV | jog | The test review was a quick jog. |
| 87 | Literal | Nominal | Motion | NOV | jump | The fence was a high jump. |
| 88 | Metaphor | Nominal | Motion | NOV | jump | The home purchase was a bungee jump. |
| 89 | Literal | Predicate | Motion | NOV | plod | The farmer plodded through the mud. |
| 90 | Metaphor | Predicate | Motion | NOV | plod | The surgeon plodded through the operation. |
| 91 | Literal | Predicate | Auditory | NOV | purr | The kitten purred on the sofa. |
| 92 | Metaphor | Predicate | Auditory | NOV | purr | The flowers purred in the sunlight. |
| 93 | Literal | Predicate | Motion | NOV | reel | The fisherman reeled in a bass. |
| 94 | Metaphor | Predicate | Motion | NOV | reel | The colonel reeled in the officers. |
| 95 | Literal | Predicate | Auditory | NOV | roar | The lion roared from his small cage. |
| 96 | Metaphor | Predicate | Auditory | NOV | roar | His curls roared amongst the bald men. |
| 97 | Literal | Nominal | Motion | NOV | roll | The bowler's throw was a straight roll. |
| 98 | Metaphor | Nominal | Motion | NOV | roll | The new roommate was a dice roll. |
| 99 | Literal | Predicate | Motion | NOV | sail | The boat sailed towards the sandy shore. |
| 100 | Metaphor | Predicate | Motion | NOV | sail | The frank speaker sailed towards a finish. |
| 101 | Literal | Predicate | Auditory | NOV | shout | Their angry coach shouted about the play. |
| 102 | Metaphor | Predicate | Auditory | NOV | shout | Her tacky shirt shouted at the interviewer. |
| 103 | Literal | Nominal | Auditory | NOV | shriek | Her contribution was a delighted shriek. |
| 104 | Metaphor | Nominal | Auditory | NOV | shriek | The purchase was a gleeful shriek. |
| 105 | Literal | Predicate | Motion | NOV | shuffle | The patient shuffled to the exit. |
| 106 | Metaphor | Predicate | Motion | NOV | shuffle | The ex-boyfriend shuffled out of her life. |
| 107 | Literal | Nominal | Motion | NOV | slither | The snake's movement was a slither. |
| 108 | Metaphor | Nominal | Motion | NOV | slither | The deal was a greedy slither. |
| 109 | Literal | Predicate | Auditory | NOV | sniff | The officer on the case sniffed at the perfume. |
| 110 | Metaphor | Predicate | Auditory | NOV | sniff | The hem of his trousers sniffed at the floor. |
| 111 | Literal | Predicate | Motion | NOV | tug | The puppy tugged at her trouser leg. |
| 112 | Metaphor | Predicate | Motion | NOV | tug | The urgent letter tugged at her sleeve. |
| 113 | Literal | Nominal | Auditory | NOV | wail | The echo was an eerie wail. |
| 114 | Metaphor | Nominal | Auditory | NOV | wail | His hangover was his liver’s wail. |
| 115 | Literal | Nominal | Motion | NOV | wander | The excursion was an afternoon wander. |
| 116 | Literal | Predicate | Motion | NOV | wander | The group wandered through the marshes. |
| 117 | Metaphor | Nominal | Motion | NOV | wander | The anthology was a literary wander. |
| 118 | Metaphor | Predicate | Motion | NOV | wander | The patient wandered through the magazine. |
| 119 | Literal | Nominal | Auditory | NOV | weep | His grief was a short weep. |
| 120 | Metaphor | Nominal | Auditory | NOV | weep | The film was a poignant weep. |

Abbreviations: No. = number, FAM = familiar, NOV = novel

**Supplemental Table 3. List of Pruned Channels Per Subject by Region of Interest (ROI)\**

| **Channel** | **Hemisphere** | **ROI** | **Participant(s) with Pruned Channel** |
| --- | --- | --- | --- |
| Ch1 | Left | Superior Frontal Gyrus (SFG) | N/A |
| Ch2 | Left | SFG | N/A |
| Ch3 | Left | Middle Frontal Gyrus (MFG) | N/A |
| Ch4 | Left | MFG | N/A |
| Ch5 | Left | Inferior Frontal Gyrus, pars triangularis (IFGtri) | P18 |
| Ch6 | Left | IFGtri | P18 |
| Ch7 | Left | IFGtri | N/A |
| Ch8 | Left | IFGtri | N/A |
| Ch9 | Left | IFGtri | N/A |
| Ch10 | Left | IFGtri | N/A |
| Ch11 | Left | IFGtri | N/A |
| Ch12 | Left | IFG, pars opercularis (IFGop) | N/A |
| Ch13 | Left | Superior Temporal Gyrus (STG) | P16 |
| Ch14 | Left | Middle Temporal Gyrus (MTG) | P16 |
| Ch15 | Left | MTG | N/A |
| Ch16 | Left | MTG | P16 |
| Ch17 | Left | Inferior Temporal Gyrus (ITG) | P18 |
| Ch18 | Left | Supramarginal Gyrus (SMG) | P8, P18 |
| Ch19 | Left | SMG | N/A |
| Ch20 | Left | SMG | N/A |
| Ch21 | Left | Angular Gyrus (AG) | P17, P18, P28 |
| Ch23 | Right | SFG | N/A |
| Ch24 | Right | MFG | N/A |
| Ch25 | Right | MFG | N/A |
| Ch26 | Right | MFG | N/A |
| Ch27 | Right | IFGtri | N/A |
| Ch28 | Right | IFGtri | P14, P20 |
| Ch29 | Right | IFGtri | N/A |
| Ch30 | Right | IFGtri | N/A |
| Ch31 | Right | IFGop | P18 |
| Ch32 | Right | IFGop | N/A |
| Ch36 | Right | STG | N/A |
| Ch37 | Right | STG | N/A |
| Ch38 | Right | STG | N/A |
| Ch39 | Right | MTG | N/A |
| Ch40 | Right | MTG | P11 |
| Ch41 | Right | MTG | P16 |
| Ch42 | Right | SMG | N/A |
| Ch43 | Right | AG | P20 |
| Ch44 | Right | AG | P20 |

Notes: Channels (Ch) are labeled with the source number followed by the number of the paired detector. Abbreviations: P = participant; N/A = not applicable (i.e., no participants with channel pruned)

**Supplemental Table 4. T-Test Outputs for the Comparison of MET > LIT and NOV/MET > FAM/MET Changes in Oxyhemoglobin per Second by Region of Interest (ROI)**

| **Time (s)** | **ROI** | **MET > LIT** | | | | **NOV/MET > FAM/MET** | | | |
| --- | --- | --- | --- | --- | --- | --- | --- | --- | --- |
|  |  | ***t*** | **df** | ***p*-value** | ***q*-value** | ***t*** | **df** | ***p*-value** | ***q*-value** |
| 0 | LSFG | 0.11 | 195 | 0.915 | 0.938 | 1.57 | 195 | 0.118 | 0.221 |
| 1 | LSFG | 1.06 | 223 | 0.289 | 0.400 | 2.24 | 223 | 0.026 | 0.070 |
| 2 | LSFG | 1.00 | 223 | 0.321 | 0.429 | 1.54 | 223 | 0.124 | 0.228 |
| 3 | LSFG | 1.10 | 195 | 0.273 | 0.382 | 0.70 | 195 | 0.483 | 0.624 |
| 4 | LSFG | 1.18 | 223 | 0.240 | 0.345 | -0.25 | 223 | 0.805 | 0.870 |
| 5 | LSFG | 1.60 | 195 | 0.111 | 0.194 | -1.44 | 195 | 0.150 | 0.263 |
| 6 | LSFG | 2.35 | 223 | 0.020 | 0.051 | -2.60 | 223 | 0.010 | 0.035* |
| 7 | LSFG | 2.98 | 223 | 0.003 | 0.011* | -3.26 | 223 | 0.001 | 0.006** |
| 8 | LSFG | 2.98 | 195 | 0.003 | 0.012* | -2.77 | 195 | 0.006 | 0.023* |
| 9 | LSFG | 3.03 | 223 | 0.003 | 0.010* | -2.37 | 223 | 0.019 | 0.055 |
| 10 | LSFG | 2.65 | 223 | 0.009 | 0.027* | -2.09 | 223 | 0.038 | 0.094 |
| 11 | LSFG | 2.00 | 195 | 0.047 | 0.102 | -2.27 | 195 | 0.024 | 0.066 |
| 12 | LSFG | 1.36 | 223 | 0.176 | 0.276 | -3.13 | 223 | 0.002 | 0.009** |
| 13 | LSFG | 0.51 | 195 | 0.610 | 0.699 | -3.51 | 195 | 0.001 | 0.003** |
| 14 | LSFG | -0.10 | 223 | 0.921 | 0.940 | -3.86 | 223 | 0.000 | 0.001** |
| 15 | LSFG | -0.48 | 223 | 0.631 | 0.717 | -3.77 | 223 | 0.000 | 0.002** |
| 16 | LSFG | -0.45 | 195 | 0.651 | 0.733 | -3.22 | 195 | 0.002 | 0.007** |
| 17 | LSFG | -0.38 | 223 | 0.706 | 0.782 | -3.11 | 223 | 0.002 | 0.009** |
| 18 | LSFG | -0.26 | 223 | 0.792 | 0.845 | -2.90 | 223 | 0.004 | 0.016* |
| 19 | LSFG | -0.17 | 195 | 0.867 | 0.904 | -2.59 | 195 | 0.010 | 0.036* |
| 20 | LSFG | 0.19 | 223 | 0.846 | 0.890 | -2.57 | 223 | 0.011 | 0.036* |
| 21 | LSFG | 0.85 | 223 | 0.398 | 0.509 | -2.16 | 223 | 0.032 | 0.082 |
| 22 | LSFG | 1.36 | 195 | 0.177 | 0.277 | -1.47 | 195 | 0.144 | 0.255 |
| 23 | LSFG | 1.57 | 223 | 0.118 | 0.203 | -0.86 | 223 | 0.393 | 0.549 |
| 24 | LSFG | 1.28 | 195 | 0.202 | 0.300 | 0.02 | 195 | 0.982 | 0.994 |
| 25 | LSFG | 1.03 | 223 | 0.302 | 0.410 | 1.02 | 223 | 0.310 | 0.458 |
| 26 | LSFG | 0.52 | 223 | 0.605 | 0.696 | 1.80 | 223 | 0.073 | 0.153 |
| 27 | LSFG | -0.52 | 195 | 0.602 | 0.696 | 2.07 | 195 | 0.040 | 0.098 |
| 28 | LSFG | -1.28 | 223 | 0.200 | 0.298 | 2.17 | 223 | 0.031 | 0.081 |
| 29 | LSFG | -2.14 | 223 | 0.034 | 0.081 | 2.05 | 223 | 0.041 | 0.100 |
| 30 | LSFG | -1.67 | 111 | 0.097 | 0.177 | 1.25 | 111 | 0.212 | 0.341 |
| 0 | LMFG | -1.51 | 195 | 0.133 | 0.224 | -2.19 | 195 | 0.029 | 0.077 |
| 1 | LMFG | -2.29 | 223 | 0.023 | 0.059 | -1.62 | 223 | 0.106 | 0.205 |
| 2 | LMFG | -2.88 | 223 | 0.004 | 0.014* | -2.88 | 223 | 0.004 | 0.017* |
| 3 | LMFG | -2.59 | 195 | 0.010 | 0.030* | -3.23 | 195 | 0.001 | 0.007** |
| 4 | LMFG | -2.58 | 223 | 0.010 | 0.031* | -4.45 | 223 | 0.000 | < 0.001*** |
| 5 | LMFG | -2.03 | 195 | 0.043 | 0.096 | -4.89 | 195 | 0.000 | < 0.001*** |
| 6 | LMFG | -1.30 | 223 | 0.195 | 0.295 | -6.37 | 223 | 0.000 | < 0.001*** |
| 7 | LMFG | -0.01 | 223 | 0.990 | 0.994 | -7.58 | 223 | 0.000 | < 0.001*** |
| 8 | LMFG | 1.29 | 195 | 0.200 | 0.298 | -7.22 | 195 | 0.000 | < 0.001*** |
| 9 | LMFG | 2.36 | 223 | 0.019 | 0.051 | -7.09 | 223 | 0.000 | < 0.001*** |
| 10 | LMFG | 2.43 | 223 | 0.016 | 0.044* | -6.77 | 223 | 0.000 | < 0.001*** |
| 11 | LMFG | 2.04 | 195 | 0.043 | 0.096 | -6.60 | 195 | 0.000 | < 0.001*** |
| 12 | LMFG | 1.72 | 223 | 0.087 | 0.161 | -7.22 | 223 | 0.000 | < 0.001*** |
| 13 | LMFG | 1.21 | 195 | 0.227 | 0.329 | -6.78 | 195 | 0.000 | < 0.001*** |
| 14 | LMFG | 0.86 | 223 | 0.392 | 0.503 | -8.15 | 223 | 0.000 | < 0.001*** |
| 15 | LMFG | 0.52 | 223 | 0.600 | 0.695 | -8.29 | 223 | 0.000 | < 0.001*** |
| 16 | LMFG | 0.45 | 195 | 0.652 | 0.733 | -7.33 | 195 | 0.000 | < 0.001*** |
| 17 | LMFG | 0.67 | 223 | 0.505 | 0.616 | -7.10 | 223 | 0.000 | < 0.001*** |
| 18 | LMFG | 1.07 | 223 | 0.287 | 0.398 | -6.46 | 223 | 0.000 | < 0.001*** |
| 19 | LMFG | 1.49 | 195 | 0.139 | 0.232 | -5.65 | 195 | 0.000 | < 0.001*** |
| 20 | LMFG | 2.13 | 223 | 0.034 | 0.082 | -5.77 | 223 | 0.000 | < 0.001*** |
| 21 | LMFG | 2.45 | 223 | 0.015 | 0.042* | -5.63 | 223 | 0.000 | < 0.001*** |
| 22 | LMFG | 2.38 | 195 | 0.019 | 0.049* | -5.09 | 195 | 0.000 | < 0.001*** |
| 23 | LMFG | 2.30 | 223 | 0.022 | 0.057 | -5.11 | 223 | 0.000 | < 0.001*** |
| 24 | LMFG | 1.82 | 195 | 0.070 | 0.138 | -4.42 | 195 | 0.000 | < 0.001*** |
| 25 | LMFG | 1.44 | 223 | 0.150 | 0.243 | -4.55 | 223 | 0.000 | < 0.001*** |
| 26 | LMFG | 1.00 | 223 | 0.321 | 0.429 | -4.34 | 223 | 0.000 | < 0.001*** |
| 27 | LMFG | 0.36 | 195 | 0.718 | 0.787 | -3.78 | 195 | 0.000 | 0.002** |
| 28 | LMFG | -0.08 | 223 | 0.936 | 0.949 | -3.94 | 223 | 0.000 | < 0.001*** |
| 29 | LMFG | -1.03 | 223 | 0.304 | 0.412 | -3.71 | 223 | 0.000 | 0.002** |
| 30 | LMFG | -1.14 | 111 | 0.258 | 0.366 | -2.62 | 111 | 0.010 | 0.035* |
| 0 | LIFGop | 0.35 | 195 | 0.725 | 0.791 | -2.11 | 195 | 0.036 | 0.092 |
| 1 | LIFGop | -0.65 | 223 | 0.514 | 0.624 | -2.02 | 223 | 0.045 | 0.106 |
| 2 | LIFGop | -0.72 | 223 | 0.474 | 0.591 | -1.38 | 223 | 0.168 | 0.287 |
| 3 | LIFGop | -1.29 | 195 | 0.200 | 0.298 | -0.94 | 195 | 0.348 | 0.500 |
| 4 | LIFGop | -1.79 | 223 | 0.075 | 0.145 | -0.77 | 223 | 0.440 | 0.598 |
| 5 | LIFGop | -1.94 | 195 | 0.053 | 0.111 | -0.68 | 195 | 0.497 | 0.630 |
| 6 | LIFGop | -1.94 | 223 | 0.053 | 0.111 | -0.96 | 223 | 0.337 | 0.487 |
| 7 | LIFGop | -1.52 | 223 | 0.130 | 0.220 | -1.35 | 223 | 0.177 | 0.294 |
| 8 | LIFGop | -0.97 | 195 | 0.333 | 0.441 | -1.57 | 195 | 0.117 | 0.221 |
| 9 | LIFGop | -0.87 | 223 | 0.385 | 0.496 | -1.74 | 223 | 0.084 | 0.174 |
| 10 | LIFGop | -0.55 | 223 | 0.585 | 0.682 | -1.58 | 223 | 0.116 | 0.221 |
| 11 | LIFGop | -0.41 | 195 | 0.684 | 0.759 | -1.19 | 195 | 0.234 | 0.371 |
| 12 | LIFGop | -0.42 | 223 | 0.674 | 0.753 | -0.74 | 223 | 0.460 | 0.607 |
| 13 | LIFGop | -0.26 | 195 | 0.794 | 0.845 | -0.07 | 195 | 0.944 | 0.966 |
| 14 | LIFGop | 0.00 | 223 | 0.996 | 0.996 | 0.46 | 223 | 0.648 | 0.752 |
| 15 | LIFGop | 0.27 | 223 | 0.788 | 0.842 | 0.88 | 223 | 0.381 | 0.535 |
| 16 | LIFGop | 0.33 | 195 | 0.744 | 0.805 | 1.01 | 195 | 0.312 | 0.459 |
| 17 | LIFGop | 0.16 | 223 | 0.873 | 0.904 | 1.13 | 223 | 0.259 | 0.399 |
| 18 | LIFGop | -0.48 | 223 | 0.631 | 0.717 | 1.07 | 223 | 0.286 | 0.429 |
| 19 | LIFGop | -1.13 | 195 | 0.261 | 0.368 | 0.97 | 195 | 0.335 | 0.485 |
| 20 | LIFGop | -2.09 | 223 | 0.038 | 0.088 | 1.10 | 223 | 0.272 | 0.414 |
| 21 | LIFGop | -2.69 | 223 | 0.008 | 0.025* | 1.36 | 223 | 0.176 | 0.294 |
| 22 | LIFGop | -2.52 | 195 | 0.013 | 0.036* | 1.57 | 195 | 0.118 | 0.221 |
| 23 | LIFGop | -2.16 | 223 | 0.032 | 0.078 | 1.98 | 223 | 0.049 | 0.112 |
| 24 | LIFGop | -1.30 | 195 | 0.197 | 0.297 | 1.99 | 195 | 0.048 | 0.112 |
| 25 | LIFGop | -0.36 | 223 | 0.717 | 0.787 | 2.45 | 223 | 0.015 | 0.047* |
| 26 | LIFGop | 0.44 | 223 | 0.663 | 0.742 | 2.71 | 223 | 0.007 | 0.026* |
| 27 | LIFGop | 1.09 | 195 | 0.277 | 0.385 | 2.80 | 195 | 0.006 | 0.022* |
| 28 | LIFGop | 1.61 | 223 | 0.108 | 0.190 | 3.45 | 223 | 0.001 | 0.004** |
| 29 | LIFGop | 2.20 | 223 | 0.029 | 0.072 | 3.82 | 223 | 0.000 | 0.001** |
| 30 | LIFGop | 1.78 | 111 | 0.077 | 0.148 | 2.38 | 111 | 0.019 | 0.056 |
| 0 | LIFGtri | 1.67 | 195 | 0.096 | 0.176 | 2.31 | 195 | 0.022 | 0.062 |
| 1 | LIFGtri | 2.40 | 223 | 0.017 | 0.047* | 3.18 | 223 | 0.002 | 0.008** |
| 2 | LIFGtri | 2.98 | 223 | 0.003 | 0.011* | 3.09 | 223 | 0.002 | 0.010* |
| 3 | LIFGtri | 2.92 | 195 | 0.004 | 0.013* | 2.42 | 195 | 0.016 | 0.049* |
| 4 | LIFGtri | 3.11 | 223 | 0.002 | 0.008** | 1.95 | 223 | 0.052 | 0.116 |
| 5 | LIFGtri | 2.95 | 195 | 0.004 | 0.012* | 1.04 | 195 | 0.301 | 0.448 |
| 6 | LIFGtri | 3.13 | 223 | 0.002 | 0.008** | 0.27 | 223 | 0.790 | 0.860 |
| 7 | LIFGtri | 3.11 | 223 | 0.002 | 0.008** | -0.70 | 223 | 0.482 | 0.624 |
| 8 | LIFGtri | 3.03 | 195 | 0.003 | 0.010* | -1.26 | 195 | 0.208 | 0.337 |
| 9 | LIFGtri | 3.57 | 223 | 0.000 | 0.002** | -1.57 | 223 | 0.118 | 0.221 |
| 10 | LIFGtri | 3.91 | 223 | 0.000 | < 0.001*** | -1.51 | 223 | 0.133 | 0.241 |
| 11 | LIFGtri | 3.57 | 195 | 0.000 | 0.002** | -1.43 | 195 | 0.155 | 0.267 |
| 12 | LIFGtri | 3.35 | 223 | 0.001 | 0.004** | -1.70 | 223 | 0.091 | 0.184 |
| 13 | LIFGtri | 2.66 | 195 | 0.008 | 0.026* | -1.84 | 195 | 0.067 | 0.144 |
| 14 | LIFGtri | 2.36 | 223 | 0.019 | 0.051 | -2.03 | 223 | 0.044 | 0.104 |
| 15 | LIFGtri | 1.86 | 223 | 0.064 | 0.129 | -2.28 | 223 | 0.024 | 0.065 |
| 16 | LIFGtri | 1.22 | 195 | 0.223 | 0.324 | -2.31 | 195 | 0.022 | 0.062 |
| 17 | LIFGtri | 0.75 | 223 | 0.454 | 0.570 | -2.60 | 223 | 0.010 | 0.035* |
| 18 | LIFGtri | 0.19 | 223 | 0.851 | 0.893 | -2.53 | 223 | 0.012 | 0.040* |
| 19 | LIFGtri | -0.18 | 195 | 0.860 | 0.899 | -2.26 | 195 | 0.025 | 0.067 |
| 20 | LIFGtri | -0.20 | 223 | 0.844 | 0.890 | -2.27 | 223 | 0.024 | 0.066 |
| 21 | LIFGtri | 0.16 | 223 | 0.871 | 0.904 | -2.10 | 223 | 0.037 | 0.093 |
| 22 | LIFGtri | 0.69 | 195 | 0.490 | 0.608 | -1.83 | 195 | 0.069 | 0.146 |
| 23 | LIFGtri | 1.35 | 223 | 0.178 | 0.278 | -1.98 | 223 | 0.049 | 0.112 |
| 24 | LIFGtri | 1.73 | 195 | 0.085 | 0.160 | -1.74 | 195 | 0.083 | 0.172 |
| 25 | LIFGtri | 1.97 | 223 | 0.050 | 0.107 | -1.47 | 223 | 0.142 | 0.252 |
| 26 | LIFGtri | 1.87 | 223 | 0.063 | 0.126 | -0.95 | 223 | 0.346 | 0.498 |
| 27 | LIFGtri | 1.43 | 195 | 0.153 | 0.247 | -0.50 | 195 | 0.618 | 0.727 |
| 28 | LIFGtri | 1.02 | 223 | 0.310 | 0.419 | 0.25 | 223 | 0.806 | 0.870 |
| 29 | LIFGtri | 0.51 | 223 | 0.613 | 0.701 | 0.61 | 223 | 0.541 | 0.669 |
| 30 | LIFGtri | 0.59 | 111 | 0.558 | 0.662 | 0.57 | 111 | 0.567 | 0.682 |
| 0 | LSTG | 0.42 | 188 | 0.677 | 0.754 | 1.86 | 188 | 0.065 | 0.139 |
| 1 | LSTG | 0.46 | 215 | 0.643 | 0.727 | 3.53 | 215 | 0.001 | 0.003** |
| 2 | LSTG | 1.04 | 215 | 0.299 | 0.409 | 4.13 | 215 | 0.000 | < 0.001*** |
| 3 | LSTG | 1.27 | 188 | 0.205 | 0.303 | 3.69 | 188 | 0.000 | 0.002** |
| 4 | LSTG | 1.75 | 215 | 0.082 | 0.154 | 3.57 | 215 | 0.000 | 0.003** |
| 5 | LSTG | 1.80 | 188 | 0.074 | 0.143 | 2.46 | 188 | 0.015 | 0.046* |
| 6 | LSTG | 2.01 | 215 | 0.046 | 0.100 | 1.43 | 215 | 0.155 | 0.267 |
| 7 | LSTG | 1.65 | 215 | 0.101 | 0.182 | 0.14 | 215 | 0.885 | 0.927 |
| 8 | LSTG | 0.69 | 188 | 0.493 | 0.610 | -0.42 | 188 | 0.677 | 0.771 |
| 9 | LSTG | -0.37 | 215 | 0.711 | 0.782 | -0.44 | 215 | 0.663 | 0.763 |
| 10 | LSTG | -1.30 | 215 | 0.194 | 0.295 | 0.00 | 215 | 0.998 | 0.998 |
| 11 | LSTG | -1.91 | 188 | 0.058 | 0.119 | 0.41 | 188 | 0.680 | 0.771 |
| 12 | LSTG | -2.62 | 215 | 0.010 | 0.029* | 1.00 | 215 | 0.317 | 0.465 |
| 13 | LSTG | -2.65 | 188 | 0.009 | 0.027* | 1.46 | 188 | 0.147 | 0.259 |
| 14 | LSTG | -2.68 | 215 | 0.008 | 0.025* | 1.69 | 215 | 0.092 | 0.184 |
| 15 | LSTG | -2.32 | 215 | 0.021 | 0.055 | 1.72 | 215 | 0.087 | 0.177 |
| 16 | LSTG | -1.92 | 188 | 0.057 | 0.116 | 1.57 | 188 | 0.119 | 0.221 |
| 17 | LSTG | -1.96 | 215 | 0.052 | 0.110 | 1.72 | 215 | 0.086 | 0.177 |
| 18 | LSTG | -1.96 | 215 | 0.052 | 0.110 | 1.99 | 215 | 0.048 | 0.112 |
| 19 | LSTG | -2.04 | 188 | 0.042 | 0.094 | 2.37 | 188 | 0.019 | 0.056 |
| 20 | LSTG | -2.47 | 215 | 0.014 | 0.041* | 3.21 | 215 | 0.002 | 0.007** |
| 21 | LSTG | -2.54 | 215 | 0.012 | 0.034* | 3.75 | 215 | 0.000 | 0.002** |
| 22 | LSTG | -2.21 | 188 | 0.029 | 0.071 | 3.75 | 188 | 0.000 | 0.002** |
| 23 | LSTG | -1.89 | 215 | 0.061 | 0.123 | 4.06 | 215 | 0.000 | < 0.001*** |
| 24 | LSTG | -1.32 | 188 | 0.188 | 0.289 | 3.76 | 188 | 0.000 | 0.002** |
| 25 | LSTG | -1.05 | 215 | 0.297 | 0.408 | 4.36 | 215 | 0.000 | < 0.001*** |
| 26 | LSTG | -0.81 | 215 | 0.419 | 0.532 | 4.80 | 215 | 0.000 | < 0.001*** |
| 27 | LSTG | -0.50 | 188 | 0.620 | 0.707 | 4.99 | 188 | 0.000 | < 0.001*** |
| 28 | LSTG | -0.36 | 215 | 0.720 | 0.787 | 5.98 | 215 | 0.000 | < 0.001*** |
| 29 | LSTG | 0.28 | 215 | 0.776 | 0.833 | 6.53 | 215 | 0.000 | < 0.001*** |
| 30 | LSTG | 0.68 | 107 | 0.496 | 0.611 | 4.62 | 107 | 0.000 | < 0.001*** |
| 0 | LMTG | 0.10 | 195 | 0.922 | 0.940 | -1.65 | 195 | 0.101 | 0.196 |
| 1 | LMTG | -0.75 | 223 | 0.456 | 0.571 | -1.69 | 223 | 0.093 | 0.185 |
| 2 | LMTG | -0.54 | 223 | 0.591 | 0.686 | -2.12 | 223 | 0.035 | 0.090 |
| 3 | LMTG | -0.41 | 195 | 0.681 | 0.757 | -2.50 | 195 | 0.013 | 0.042* |
| 4 | LMTG | -0.16 | 223 | 0.870 | 0.904 | -3.02 | 223 | 0.003 | 0.012* |
| 5 | LMTG | -0.10 | 195 | 0.921 | 0.940 | -3.03 | 195 | 0.003 | 0.012* |
| 6 | LMTG | -0.18 | 223 | 0.857 | 0.898 | -3.46 | 223 | 0.001 | 0.004** |
| 7 | LMTG | -0.60 | 223 | 0.551 | 0.657 | -3.62 | 223 | 0.000 | 0.002** |
| 8 | LMTG | -0.95 | 195 | 0.342 | 0.449 | -3.33 | 195 | 0.001 | 0.006** |
| 9 | LMTG | -1.43 | 223 | 0.153 | 0.247 | -3.38 | 223 | 0.001 | 0.005** |
| 10 | LMTG | -1.79 | 223 | 0.074 | 0.144 | -3.21 | 223 | 0.002 | 0.007** |
| 11 | LMTG | -1.95 | 195 | 0.053 | 0.111 | -2.91 | 195 | 0.004 | 0.016* |
| 12 | LMTG | -2.32 | 223 | 0.021 | 0.055 | -2.95 | 223 | 0.004 | 0.015* |
| 13 | LMTG | -2.36 | 195 | 0.019 | 0.051 | -2.51 | 195 | 0.013 | 0.042* |
| 14 | LMTG | -2.68 | 223 | 0.008 | 0.025* | -2.57 | 223 | 0.011 | 0.036* |
| 15 | LMTG | -2.96 | 223 | 0.003 | 0.012* | -2.44 | 223 | 0.015 | 0.047* |
| 16 | LMTG | -3.08 | 195 | 0.002 | 0.009** | -2.26 | 195 | 0.025 | 0.067 |
| 17 | LMTG | -3.63 | 223 | 0.000 | 0.002** | -2.28 | 223 | 0.023 | 0.065 |
| 18 | LMTG | -3.84 | 223 | 0.000 | < 0.001*** | -2.00 | 223 | 0.047 | 0.111 |
| 19 | LMTG | -3.57 | 195 | 0.000 | 0.002** | -1.56 | 195 | 0.120 | 0.222 |
| 20 | LMTG | -3.59 | 223 | 0.000 | 0.002** | -1.37 | 223 | 0.171 | 0.289 |
| 21 | LMTG | -3.06 | 223 | 0.002 | 0.009** | -1.02 | 223 | 0.308 | 0.457 |
| 22 | LMTG | -2.10 | 195 | 0.037 | 0.087 | -0.59 | 195 | 0.553 | 0.674 |
| 23 | LMTG | -1.36 | 223 | 0.174 | 0.274 | -0.07 | 223 | 0.943 | 0.966 |
| 24 | LMTG | -0.59 | 195 | 0.559 | 0.662 | 0.50 | 195 | 0.615 | 0.725 |
| 25 | LMTG | -0.03 | 223 | 0.978 | 0.984 | 1.22 | 223 | 0.222 | 0.355 |
| 26 | LMTG | 0.25 | 223 | 0.800 | 0.848 | 1.87 | 223 | 0.063 | 0.135 |
| 27 | LMTG | 0.31 | 195 | 0.755 | 0.814 | 2.47 | 195 | 0.015 | 0.046* |
| 28 | LMTG | 0.34 | 223 | 0.734 | 0.799 | 2.97 | 223 | 0.003 | 0.014* |
| 29 | LMTG | 0.77 | 223 | 0.445 | 0.560 | 3.18 | 223 | 0.002 | 0.008** |
| 30 | LMTG | 0.88 | 111 | 0.379 | 0.491 | 1.89 | 111 | 0.062 | 0.134 |
| 0 | LITG | 2.18 | 188 | 0.031 | 0.075 | -4.79 | 188 | 0.000 | < 0.001*** |
| 1 | LITG | 0.97 | 215 | 0.331 | 0.440 | -5.08 | 215 | 0.000 | < 0.001*** |
| 2 | LITG | 1.44 | 215 | 0.150 | 0.243 | -5.08 | 215 | 0.000 | < 0.001*** |
| 3 | LITG | 1.49 | 188 | 0.138 | 0.231 | -4.67 | 188 | 0.000 | < 0.001*** |
| 4 | LITG | 1.97 | 215 | 0.051 | 0.109 | -5.00 | 215 | 0.000 | < 0.001*** |
| 5 | LITG | 2.10 | 188 | 0.037 | 0.086 | -5.03 | 188 | 0.000 | < 0.001*** |
| 6 | LITG | 2.80 | 215 | 0.006 | 0.018* | -5.38 | 215 | 0.000 | < 0.001*** |
| 7 | LITG | 3.29 | 215 | 0.001 | 0.005** | -5.13 | 215 | 0.000 | < 0.001*** |
| 8 | LITG | 3.44 | 188 | 0.001 | 0.003** | -4.29 | 188 | 0.000 | < 0.001*** |
| 9 | LITG | 3.96 | 215 | 0.000 | < 0.001*** | -3.94 | 215 | 0.000 | < 0.001*** |
| 10 | LITG | 4.22 | 215 | 0.000 | < 0.001*** | -3.23 | 215 | 0.001 | 0.007** |
| 11 | LITG | 4.10 | 188 | 0.000 | < 0.001*** | -2.58 | 188 | 0.011 | 0.036* |
| 12 | LITG | 4.38 | 215 | 0.000 | < 0.001*** | -2.48 | 215 | 0.014 | 0.045* |
| 13 | LITG | 3.98 | 188 | 0.000 | < 0.001*** | -2.07 | 188 | 0.040 | 0.098 |
| 14 | LITG | 4.14 | 215 | 0.000 | < 0.001*** | -1.89 | 215 | 0.061 | 0.133 |
| 15 | LITG | 4.14 | 215 | 0.000 | < 0.001*** | -1.49 | 215 | 0.138 | 0.247 |
| 16 | LITG | 3.92 | 188 | 0.000 | < 0.001*** | -1.02 | 188 | 0.310 | 0.458 |
| 17 | LITG | 4.37 | 215 | 0.000 | < 0.001*** | -0.71 | 215 | 0.478 | 0.620 |
| 18 | LITG | 4.61 | 215 | 0.000 | < 0.001*** | -0.59 | 215 | 0.553 | 0.674 |
| 19 | LITG | 4.36 | 188 | 0.000 | < 0.001*** | -0.67 | 188 | 0.501 | 0.633 |
| 20 | LITG | 4.42 | 215 | 0.000 | < 0.001*** | -0.99 | 215 | 0.323 | 0.472 |
| 21 | LITG | 4.12 | 215 | 0.000 | < 0.001*** | -1.26 | 215 | 0.210 | 0.339 |
| 22 | LITG | 3.68 | 188 | 0.000 | 0.001** | -1.38 | 188 | 0.171 | 0.289 |
| 23 | LITG | 3.97 | 215 | 0.000 | < 0.001*** | -1.51 | 215 | 0.133 | 0.241 |
| 24 | LITG | 3.86 | 188 | 0.000 | < 0.001*** | -1.23 | 188 | 0.222 | 0.355 |
| 25 | LITG | 4.61 | 215 | 0.000 | < 0.001*** | -0.75 | 215 | 0.455 | 0.606 |
| 26 | LITG | 5.20 | 215 | 0.000 | < 0.001*** | -0.27 | 215 | 0.785 | 0.860 |
| 27 | LITG | 5.50 | 188 | 0.000 | < 0.001*** | 0.08 | 188 | 0.933 | 0.959 |
| 28 | LITG | 6.05 | 215 | 0.000 | < 0.001*** | -0.16 | 215 | 0.877 | 0.924 |
| 29 | LITG | 5.96 | 215 | 0.000 | < 0.001*** | 0.01 | 215 | 0.989 | 0.996 |
| 30 | LITG | 4.01 | 107 | 0.000 | < 0.001*** | -0.32 | 107 | 0.747 | 0.834 |
| 0 | LSMG | 3.93 | 195 | 0.000 | < 0.001*** | 0.29 | 195 | 0.774 | 0.857 |
| 1 | LSMG | 2.71 | 223 | 0.007 | 0.023* | 0.12 | 223 | 0.908 | 0.942 |
| 2 | LSMG | 2.74 | 223 | 0.007 | 0.022* | -0.93 | 223 | 0.352 | 0.503 |
| 3 | LSMG | 2.34 | 195 | 0.020 | 0.053 | -1.32 | 195 | 0.188 | 0.309 |
| 4 | LSMG | 2.64 | 223 | 0.009 | 0.027* | -2.10 | 223 | 0.037 | 0.093 |
| 5 | LSMG | 2.42 | 195 | 0.017 | 0.046* | -2.85 | 195 | 0.005 | 0.019* |
| 6 | LSMG | 2.72 | 223 | 0.007 | 0.023* | -3.90 | 223 | 0.000 | 0.001** |
| 7 | LSMG | 2.91 | 223 | 0.004 | 0.013* | -4.77 | 223 | 0.000 | < 0.001*** |
| 8 | LSMG | 2.95 | 195 | 0.004 | 0.012* | -4.71 | 195 | 0.000 | < 0.001*** |
| 9 | LSMG | 3.32 | 223 | 0.001 | 0.004** | -4.17 | 223 | 0.000 | < 0.001*** |
| 10 | LSMG | 3.65 | 223 | 0.000 | 0.001** | -2.34 | 223 | 0.020 | 0.058 |
| 11 | LSMG | 3.72 | 195 | 0.000 | 0.001** | -0.81 | 195 | 0.418 | 0.575 |
| 12 | LSMG | 4.11 | 223 | 0.000 | < 0.001*** | 0.15 | 223 | 0.884 | 0.927 |
| 13 | LSMG | 3.92 | 195 | 0.000 | < 0.001*** | 0.60 | 195 | 0.551 | 0.674 |
| 14 | LSMG | 4.26 | 223 | 0.000 | < 0.001*** | 0.19 | 223 | 0.852 | 0.916 |
| 15 | LSMG | 4.37 | 223 | 0.000 | < 0.001*** | -0.65 | 223 | 0.517 | 0.644 |
| 16 | LSMG | 4.29 | 195 | 0.000 | < 0.001*** | -1.31 | 195 | 0.191 | 0.313 |
| 17 | LSMG | 4.76 | 223 | 0.000 | < 0.001*** | -2.02 | 223 | 0.045 | 0.106 |
| 18 | LSMG | 4.79 | 223 | 0.000 | < 0.001*** | -2.40 | 223 | 0.017 | 0.051 |
| 19 | LSMG | 4.51 | 195 | 0.000 | < 0.001*** | -2.27 | 195 | 0.024 | 0.066 |
| 20 | LSMG | 4.84 | 223 | 0.000 | < 0.001*** | -2.21 | 223 | 0.028 | 0.074 |
| 21 | LSMG | 5.04 | 223 | 0.000 | < 0.001*** | -2.05 | 223 | 0.041 | 0.100 |
| 22 | LSMG | 5.00 | 195 | 0.000 | < 0.001*** | -1.69 | 195 | 0.092 | 0.184 |
| 23 | LSMG | 5.70 | 223 | 0.000 | < 0.001*** | -1.68 | 223 | 0.095 | 0.188 |
| 24 | LSMG | 5.36 | 195 | 0.000 | < 0.001*** | -1.55 | 195 | 0.124 | 0.228 |
| 25 | LSMG | 5.98 | 223 | 0.000 | < 0.001*** | -1.53 | 223 | 0.129 | 0.235 |
| 26 | LSMG | 6.07 | 223 | 0.000 | < 0.001*** | -1.38 | 223 | 0.170 | 0.289 |
| 27 | LSMG | 5.95 | 195 | 0.000 | < 0.001*** | -1.18 | 195 | 0.238 | 0.377 |
| 28 | LSMG | 6.31 | 223 | 0.000 | < 0.001*** | -1.22 | 223 | 0.223 | 0.355 |
| 29 | LSMG | 6.85 | 223 | 0.000 | < 0.001*** | -1.53 | 223 | 0.128 | 0.234 |
| 30 | LSMG | 4.72 | 111 | 0.000 | < 0.001*** | -1.44 | 111 | 0.154 | 0.267 |
| 0 | LAG | 0.13 | 174 | 0.898 | 0.924 | 1.13 | 174 | 0.258 | 0.399 |
| 1 | LAG | -1.32 | 199 | 0.190 | 0.290 | 0.92 | 199 | 0.358 | 0.510 |
| 2 | LAG | -0.71 | 199 | 0.479 | 0.595 | 0.27 | 199 | 0.786 | 0.860 |
| 3 | LAG | -0.55 | 174 | 0.582 | 0.681 | 0.00 | 174 | 0.998 | 0.998 |
| 4 | LAG | 0.08 | 199 | 0.933 | 0.947 | -0.27 | 199 | 0.788 | 0.860 |
| 5 | LAG | 0.51 | 174 | 0.608 | 0.698 | -0.45 | 174 | 0.651 | 0.754 |
| 6 | LAG | 1.29 | 199 | 0.197 | 0.297 | -0.44 | 199 | 0.659 | 0.762 |
| 7 | LAG | 2.05 | 199 | 0.041 | 0.094 | -0.16 | 199 | 0.872 | 0.924 |
| 8 | LAG | 2.64 | 174 | 0.009 | 0.028* | 0.15 | 174 | 0.883 | 0.927 |
| 9 | LAG | 3.57 | 199 | 0.000 | 0.002** | 0.48 | 199 | 0.630 | 0.737 |
| 10 | LAG | 4.21 | 199 | 0.000 | < 0.001*** | 0.75 | 199 | 0.457 | 0.607 |
| 11 | LAG | 4.29 | 174 | 0.000 | < 0.001*** | 0.56 | 174 | 0.574 | 0.686 |
| 12 | LAG | 4.83 | 199 | 0.000 | < 0.001*** | -0.16 | 199 | 0.873 | 0.924 |
| 13 | LAG | 4.86 | 174 | 0.000 | < 0.001*** | -0.99 | 174 | 0.323 | 0.472 |
| 14 | LAG | 5.70 | 199 | 0.000 | < 0.001*** | -1.17 | 199 | 0.242 | 0.381 |
| 15 | LAG | 6.18 | 199 | 0.000 | < 0.001*** | -1.15 | 199 | 0.250 | 0.389 |
| 16 | LAG | 6.00 | 174 | 0.000 | < 0.001*** | -0.82 | 174 | 0.414 | 0.571 |
| 17 | LAG | 6.25 | 199 | 0.000 | < 0.001*** | -0.58 | 199 | 0.565 | 0.682 |
| 18 | LAG | 5.67 | 199 | 0.000 | < 0.001*** | -0.36 | 199 | 0.722 | 0.815 |
| 19 | LAG | 4.61 | 174 | 0.000 | < 0.001*** | -0.26 | 174 | 0.795 | 0.864 |
| 20 | LAG | 4.45 | 199 | 0.000 | < 0.001*** | -0.33 | 199 | 0.740 | 0.830 |
| 21 | LAG | 4.41 | 199 | 0.000 | < 0.001*** | -0.44 | 199 | 0.661 | 0.762 |
| 22 | LAG | 4.44 | 174 | 0.000 | < 0.001*** | -0.56 | 174 | 0.576 | 0.687 |
| 23 | LAG | 5.10 | 199 | 0.000 | < 0.001*** | -0.75 | 199 | 0.451 | 0.605 |
| 24 | LAG | 4.79 | 174 | 0.000 | < 0.001*** | -0.73 | 174 | 0.466 | 0.611 |
| 25 | LAG | 5.03 | 199 | 0.000 | < 0.001*** | -0.41 | 199 | 0.680 | 0.771 |
| 26 | LAG | 4.70 | 199 | 0.000 | < 0.001*** | -0.16 | 199 | 0.872 | 0.924 |
| 27 | LAG | 4.14 | 174 | 0.000 | < 0.001*** | -0.14 | 174 | 0.888 | 0.929 |
| 28 | LAG | 4.27 | 199 | 0.000 | < 0.001*** | -0.58 | 199 | 0.562 | 0.681 |
| 29 | LAG | 4.63 | 199 | 0.000 | < 0.001*** | -0.59 | 199 | 0.554 | 0.674 |
| 30 | LAG | 3.38 | 99 | 0.001 | 0.004** | -0.43 | 99 | 0.667 | 0.764 |
| 0 | RSFG | 0.30 | 195 | 0.763 | 0.821 | 0.01 | 195 | 0.992 | 0.996 |
| 1 | RSFG | 1.46 | 223 | 0.145 | 0.239 | 1.48 | 223 | 0.139 | 0.248 |
| 2 | RSFG | -0.33 | 223 | 0.744 | 0.805 | 1.37 | 223 | 0.173 | 0.289 |
| 3 | RSFG | -1.52 | 195 | 0.131 | 0.221 | 1.10 | 195 | 0.271 | 0.414 |
| 4 | RSFG | -2.55 | 223 | 0.012 | 0.033* | 0.76 | 223 | 0.448 | 0.602 |
| 5 | RSFG | -2.18 | 195 | 0.030 | 0.075 | -0.02 | 195 | 0.985 | 0.994 |
| 6 | RSFG | -1.91 | 223 | 0.058 | 0.118 | -1.08 | 223 | 0.283 | 0.426 |
| 7 | RSFG | -1.43 | 223 | 0.154 | 0.247 | -2.33 | 223 | 0.021 | 0.059 |
| 8 | RSFG | -1.23 | 195 | 0.219 | 0.319 | -2.98 | 195 | 0.003 | 0.014* |
| 9 | RSFG | -1.60 | 223 | 0.112 | 0.195 | -3.44 | 223 | 0.001 | 0.004** |
| 10 | RSFG | -2.40 | 223 | 0.017 | 0.047* | -3.25 | 223 | 0.001 | 0.007** |
| 11 | RSFG | -3.00 | 195 | 0.003 | 0.011* | -2.79 | 195 | 0.006 | 0.022* |
| 12 | RSFG | -3.80 | 223 | 0.000 | < 0.001*** | -2.77 | 223 | 0.006 | 0.023* |
| 13 | RSFG | -3.59 | 195 | 0.000 | 0.002** | -2.32 | 195 | 0.021 | 0.060 |
| 14 | RSFG | -3.42 | 223 | 0.001 | 0.003** | -1.58 | 223 | 0.115 | 0.220 |
| 15 | RSFG | -2.95 | 223 | 0.004 | 0.012* | -0.97 | 223 | 0.332 | 0.482 |
| 16 | RSFG | -2.33 | 195 | 0.021 | 0.054 | -0.46 | 195 | 0.647 | 0.752 |
| 17 | RSFG | -2.14 | 223 | 0.033 | 0.080 | -0.14 | 223 | 0.893 | 0.930 |
| 18 | RSFG | -1.99 | 223 | 0.048 | 0.104 | 0.27 | 223 | 0.789 | 0.860 |
| 19 | RSFG | -1.90 | 195 | 0.059 | 0.119 | 0.49 | 195 | 0.622 | 0.730 |
| 20 | RSFG | -2.06 | 223 | 0.040 | 0.091 | 0.75 | 223 | 0.455 | 0.606 |
| 21 | RSFG | -1.88 | 223 | 0.061 | 0.123 | 0.73 | 223 | 0.469 | 0.612 |
| 22 | RSFG | -1.52 | 195 | 0.130 | 0.220 | 0.66 | 195 | 0.513 | 0.644 |
| 23 | RSFG | -1.33 | 223 | 0.185 | 0.286 | 0.64 | 223 | 0.523 | 0.650 |
| 24 | RSFG | -1.16 | 195 | 0.247 | 0.353 | 0.42 | 195 | 0.675 | 0.770 |
| 25 | RSFG | -1.12 | 223 | 0.265 | 0.372 | 0.31 | 223 | 0.760 | 0.845 |
| 26 | RSFG | -1.13 | 223 | 0.259 | 0.366 | 0.09 | 223 | 0.932 | 0.959 |
| 27 | RSFG | -1.17 | 195 | 0.245 | 0.351 | 0.16 | 195 | 0.873 | 0.924 |
| 28 | RSFG | -1.44 | 223 | 0.150 | 0.243 | 0.09 | 223 | 0.927 | 0.959 |
| 29 | RSFG | -1.81 | 223 | 0.072 | 0.141 | 0.57 | 223 | 0.572 | 0.685 |
| 30 | RSFG | -1.69 | 111 | 0.094 | 0.173 | 0.43 | 111 | 0.665 | 0.764 |
| 0 | RMFG | -2.67 | 195 | 0.008 | 0.026* | -1.25 | 195 | 0.211 | 0.340 |
| 1 | RMFG | -2.82 | 223 | 0.005 | 0.017* | 0.84 | 223 | 0.404 | 0.560 |
| 2 | RMFG | -3.67 | 223 | 0.000 | 0.001** | -0.07 | 223 | 0.947 | 0.967 |
| 3 | RMFG | -3.44 | 195 | 0.001 | 0.003** | -0.28 | 195 | 0.783 | 0.860 |
| 4 | RMFG | -3.79 | 223 | 0.000 | < 0.001*** | -1.05 | 223 | 0.294 | 0.439 |
| 5 | RMFG | -3.12 | 195 | 0.002 | 0.008** | -2.03 | 195 | 0.044 | 0.105 |
| 6 | RMFG | -2.51 | 223 | 0.013 | 0.036* | -3.55 | 223 | 0.000 | 0.003** |
| 7 | RMFG | -1.75 | 223 | 0.082 | 0.154 | -5.16 | 223 | 0.000 | < 0.001*** |
| 8 | RMFG | -1.34 | 195 | 0.182 | 0.282 | -5.86 | 195 | 0.000 | < 0.001*** |
| 9 | RMFG | -1.48 | 223 | 0.141 | 0.234 | -6.60 | 223 | 0.000 | < 0.001*** |
| 10 | RMFG | -1.69 | 223 | 0.093 | 0.172 | -6.00 | 223 | 0.000 | < 0.001*** |
| 11 | RMFG | -1.65 | 195 | 0.100 | 0.181 | -5.03 | 195 | 0.000 | < 0.001*** |
| 12 | RMFG | -1.85 | 223 | 0.066 | 0.132 | -5.00 | 223 | 0.000 | < 0.001*** |
| 13 | RMFG | -1.54 | 195 | 0.125 | 0.213 | -4.54 | 195 | 0.000 | < 0.001*** |
| 14 | RMFG | -1.37 | 223 | 0.173 | 0.273 | -4.46 | 223 | 0.000 | < 0.001*** |
| 15 | RMFG | -0.97 | 223 | 0.334 | 0.441 | -4.37 | 223 | 0.000 | < 0.001*** |
| 16 | RMFG | -0.37 | 195 | 0.708 | 0.782 | -3.93 | 195 | 0.000 | < 0.001*** |
| 17 | RMFG | 0.13 | 223 | 0.896 | 0.924 | -3.96 | 223 | 0.000 | < 0.001*** |
| 18 | RMFG | 0.58 | 223 | 0.561 | 0.663 | -3.44 | 223 | 0.001 | 0.004** |
| 19 | RMFG | 0.88 | 195 | 0.381 | 0.492 | -2.73 | 195 | 0.007 | 0.026* |
| 20 | RMFG | 1.35 | 223 | 0.179 | 0.278 | -2.42 | 223 | 0.016 | 0.049* |
| 21 | RMFG | 1.58 | 223 | 0.116 | 0.200 | -2.10 | 223 | 0.037 | 0.093 |
| 22 | RMFG | 1.60 | 195 | 0.111 | 0.194 | -1.83 | 195 | 0.069 | 0.146 |
| 23 | RMFG | 1.54 | 223 | 0.124 | 0.212 | -1.98 | 223 | 0.049 | 0.112 |
| 24 | RMFG | 1.11 | 195 | 0.270 | 0.378 | -1.73 | 195 | 0.086 | 0.176 |
| 25 | RMFG | 0.67 | 223 | 0.505 | 0.616 | -1.96 | 223 | 0.051 | 0.114 |
| 26 | RMFG | -0.08 | 223 | 0.933 | 0.947 | -2.26 | 223 | 0.025 | 0.067 |
| 27 | RMFG | -1.25 | 195 | 0.212 | 0.310 | -2.19 | 195 | 0.030 | 0.078 |
| 28 | RMFG | -2.43 | 223 | 0.016 | 0.044* | -2.46 | 223 | 0.015 | 0.046* |
| 29 | RMFG | -3.75 | 223 | 0.000 | 0.001** | -2.31 | 223 | 0.022 | 0.062 |
| 30 | RMFG | -3.00 | 111 | 0.003 | 0.012* | -1.82 | 111 | 0.071 | 0.150 |
| 0 | RIFGop | 0.24 | 195 | 0.813 | 0.859 | -1.98 | 195 | 0.050 | 0.112 |
| 1 | RIFGop | -0.97 | 223 | 0.333 | 0.441 | 0.01 | 223 | 0.991 | 0.996 |
| 2 | RIFGop | -0.67 | 223 | 0.504 | 0.616 | 0.68 | 223 | 0.496 | 0.630 |
| 3 | RIFGop | -0.83 | 195 | 0.406 | 0.517 | 1.17 | 195 | 0.245 | 0.383 |
| 4 | RIFGop | -0.54 | 223 | 0.591 | 0.686 | 1.39 | 223 | 0.166 | 0.284 |
| 5 | RIFGop | -0.52 | 195 | 0.605 | 0.696 | 1.08 | 195 | 0.282 | 0.426 |
| 6 | RIFGop | -0.67 | 223 | 0.504 | 0.616 | 0.77 | 223 | 0.439 | 0.598 |
| 7 | RIFGop | -0.86 | 223 | 0.390 | 0.501 | 0.31 | 223 | 0.760 | 0.845 |
| 8 | RIFGop | -0.96 | 195 | 0.339 | 0.447 | -0.28 | 195 | 0.780 | 0.860 |
| 9 | RIFGop | -1.29 | 223 | 0.198 | 0.297 | -0.88 | 223 | 0.377 | 0.534 |
| 10 | RIFGop | -1.40 | 223 | 0.162 | 0.259 | -0.83 | 223 | 0.406 | 0.562 |
| 11 | RIFGop | -1.09 | 195 | 0.275 | 0.383 | -0.28 | 195 | 0.776 | 0.857 |
| 12 | RIFGop | -0.37 | 223 | 0.709 | 0.782 | 0.18 | 223 | 0.857 | 0.920 |
| 13 | RIFGop | 0.78 | 195 | 0.434 | 0.550 | 0.72 | 195 | 0.473 | 0.615 |
| 14 | RIFGop | 1.65 | 223 | 0.101 | 0.182 | 1.99 | 223 | 0.048 | 0.112 |
| 15 | RIFGop | 2.03 | 223 | 0.044 | 0.097 | 2.94 | 223 | 0.004 | 0.015* |
| 16 | RIFGop | 2.31 | 195 | 0.022 | 0.057 | 3.28 | 195 | 0.001 | 0.006** |
| 17 | RIFGop | 3.02 | 223 | 0.003 | 0.010* | 3.43 | 223 | 0.001 | 0.004** |
| 18 | RIFGop | 3.47 | 223 | 0.001 | 0.003** | 2.91 | 223 | 0.004 | 0.016* |
| 19 | RIFGop | 3.72 | 195 | 0.000 | 0.001** | 2.10 | 195 | 0.037 | 0.093 |
| 20 | RIFGop | 4.52 | 223 | 0.000 | < 0.001*** | 1.80 | 223 | 0.073 | 0.153 |
| 21 | RIFGop | 5.00 | 223 | 0.000 | < 0.001*** | 1.71 | 223 | 0.088 | 0.179 |
| 22 | RIFGop | 5.07 | 195 | 0.000 | < 0.001*** | 1.72 | 195 | 0.088 | 0.178 |
| 23 | RIFGop | 5.65 | 223 | 0.000 | < 0.001*** | 2.06 | 223 | 0.040 | 0.098 |
| 24 | RIFGop | 5.32 | 195 | 0.000 | < 0.001*** | 2.54 | 195 | 0.012 | 0.039* |
| 25 | RIFGop | 5.64 | 223 | 0.000 | < 0.001*** | 3.49 | 223 | 0.001 | 0.004** |
| 26 | RIFGop | 5.57 | 223 | 0.000 | < 0.001*** | 3.90 | 223 | 0.000 | 0.001** |
| 27 | RIFGop | 4.72 | 195 | 0.000 | < 0.001*** | 3.43 | 195 | 0.001 | 0.004** |
| 28 | RIFGop | 4.26 | 223 | 0.000 | < 0.001*** | 3.33 | 223 | 0.001 | 0.005** |
| 29 | RIFGop | 3.51 | 223 | 0.001 | 0.002** | 2.79 | 223 | 0.006 | 0.022* |
| 30 | RIFGop | 2.58 | 111 | 0.011 | 0.032* | 1.67 | 111 | 0.098 | 0.191 |
| 0 | RIFGtri | -0.03 | 195 | 0.977 | 0.984 | -1.33 | 195 | 0.186 | 0.306 |
| 1 | RIFGtri | 0.64 | 223 | 0.523 | 0.632 | -0.32 | 223 | 0.746 | 0.834 |
| 2 | RIFGtri | 1.71 | 223 | 0.089 | 0.164 | 0.05 | 223 | 0.961 | 0.978 |
| 3 | RIFGtri | 2.36 | 195 | 0.019 | 0.051 | 0.61 | 195 | 0.543 | 0.670 |
| 4 | RIFGtri | 2.95 | 223 | 0.003 | 0.012* | 0.65 | 223 | 0.517 | 0.644 |
| 5 | RIFGtri | 2.61 | 195 | 0.010 | 0.029* | -0.03 | 195 | 0.978 | 0.993 |
| 6 | RIFGtri | 2.50 | 223 | 0.013 | 0.037* | -1.33 | 223 | 0.185 | 0.306 |
| 7 | RIFGtri | 2.10 | 223 | 0.037 | 0.086 | -3.02 | 223 | 0.003 | 0.012* |
| 8 | RIFGtri | 1.46 | 195 | 0.145 | 0.239 | -4.21 | 195 | 0.000 | < 0.001*** |
| 9 | RIFGtri | 0.98 | 223 | 0.330 | 0.440 | -5.37 | 223 | 0.000 | < 0.001*** |
| 10 | RIFGtri | 0.78 | 223 | 0.437 | 0.552 | -5.19 | 223 | 0.000 | < 0.001*** |
| 11 | RIFGtri | 0.72 | 195 | 0.472 | 0.589 | -4.05 | 195 | 0.000 | < 0.001*** |
| 12 | RIFGtri | 0.94 | 223 | 0.348 | 0.456 | -3.41 | 223 | 0.001 | 0.004** |
| 13 | RIFGtri | 1.24 | 195 | 0.215 | 0.314 | -2.56 | 195 | 0.011 | 0.037* |
| 14 | RIFGtri | 1.66 | 223 | 0.098 | 0.177 | -2.10 | 223 | 0.037 | 0.093 |
| 15 | RIFGtri | 1.86 | 223 | 0.064 | 0.129 | -2.04 | 223 | 0.042 | 0.102 |
| 16 | RIFGtri | 1.78 | 195 | 0.077 | 0.148 | -2.15 | 195 | 0.033 | 0.085 |
| 17 | RIFGtri | 1.82 | 223 | 0.070 | 0.138 | -2.53 | 223 | 0.012 | 0.040* |
| 18 | RIFGtri | 1.60 | 223 | 0.112 | 0.195 | -2.41 | 223 | 0.017 | 0.051 |
| 19 | RIFGtri | 1.28 | 195 | 0.204 | 0.302 | -1.95 | 195 | 0.053 | 0.116 |
| 20 | RIFGtri | 1.33 | 223 | 0.186 | 0.287 | -1.65 | 223 | 0.101 | 0.196 |
| 21 | RIFGtri | 1.40 | 223 | 0.164 | 0.261 | -1.16 | 223 | 0.246 | 0.384 |
| 22 | RIFGtri | 1.59 | 195 | 0.113 | 0.196 | -0.74 | 195 | 0.461 | 0.607 |
| 23 | RIFGtri | 1.94 | 223 | 0.054 | 0.112 | -0.61 | 223 | 0.545 | 0.671 |
| 24 | RIFGtri | 1.93 | 195 | 0.055 | 0.114 | -0.48 | 195 | 0.631 | 0.737 |
| 25 | RIFGtri | 1.95 | 223 | 0.053 | 0.111 | -0.69 | 223 | 0.489 | 0.629 |
| 26 | RIFGtri | 1.77 | 223 | 0.079 | 0.150 | -1.12 | 223 | 0.264 | 0.406 |
| 27 | RIFGtri | 1.19 | 195 | 0.234 | 0.337 | -1.37 | 195 | 0.173 | 0.289 |
| 28 | RIFGtri | 0.93 | 223 | 0.355 | 0.464 | -1.27 | 223 | 0.206 | 0.336 |
| 29 | RIFGtri | 0.31 | 223 | 0.755 | 0.814 | -0.69 | 223 | 0.489 | 0.629 |
| 30 | RIFGtri | 0.46 | 111 | 0.644 | 0.727 | -0.43 | 111 | 0.670 | 0.766 |
| 0 | RSTG | -0.56 | 195 | 0.577 | 0.679 | 1.08 | 195 | 0.283 | 0.426 |
| 1 | RSTG | -0.62 | 223 | 0.533 | 0.641 | 0.77 | 223 | 0.443 | 0.599 |
| 2 | RSTG | -0.58 | 223 | 0.562 | 0.663 | 0.14 | 223 | 0.890 | 0.929 |
| 3 | RSTG | -0.55 | 195 | 0.583 | 0.681 | -0.19 | 195 | 0.846 | 0.912 |
| 4 | RSTG | -0.59 | 223 | 0.556 | 0.661 | -0.74 | 223 | 0.463 | 0.608 |
| 5 | RSTG | -0.56 | 195 | 0.579 | 0.680 | -1.50 | 195 | 0.134 | 0.242 |
| 6 | RSTG | -0.69 | 223 | 0.494 | 0.610 | -2.28 | 223 | 0.023 | 0.065 |
| 7 | RSTG | -0.90 | 223 | 0.367 | 0.479 | -2.68 | 223 | 0.008 | 0.029* |
| 8 | RSTG | -1.27 | 195 | 0.206 | 0.303 | -2.53 | 195 | 0.012 | 0.040* |
| 9 | RSTG | -1.83 | 223 | 0.068 | 0.135 | -2.50 | 223 | 0.013 | 0.042* |
| 10 | RSTG | -2.13 | 223 | 0.034 | 0.082 | -1.99 | 223 | 0.048 | 0.112 |
| 11 | RSTG | -2.09 | 195 | 0.038 | 0.087 | -1.37 | 195 | 0.172 | 0.289 |
| 12 | RSTG | -2.05 | 223 | 0.042 | 0.094 | -1.17 | 223 | 0.242 | 0.381 |
| 13 | RSTG | -1.31 | 195 | 0.192 | 0.292 | -0.92 | 195 | 0.360 | 0.511 |
| 14 | RSTG | -0.45 | 223 | 0.656 | 0.736 | -0.51 | 223 | 0.612 | 0.723 |
| 15 | RSTG | 0.63 | 223 | 0.528 | 0.637 | -0.26 | 223 | 0.799 | 0.866 |
| 16 | RSTG | 1.47 | 195 | 0.144 | 0.239 | -0.09 | 195 | 0.929 | 0.959 |
| 17 | RSTG | 2.08 | 223 | 0.039 | 0.089 | -0.17 | 223 | 0.862 | 0.921 |
| 18 | RSTG | 2.04 | 223 | 0.042 | 0.094 | -0.37 | 223 | 0.709 | 0.802 |
| 19 | RSTG | 1.49 | 195 | 0.138 | 0.231 | -0.53 | 195 | 0.597 | 0.710 |
| 20 | RSTG | 0.85 | 223 | 0.399 | 0.509 | -0.68 | 223 | 0.495 | 0.630 |
| 21 | RSTG | -0.03 | 223 | 0.975 | 0.984 | -0.85 | 223 | 0.396 | 0.551 |
| 22 | RSTG | -0.62 | 195 | 0.539 | 0.647 | -0.87 | 195 | 0.384 | 0.538 |
| 23 | RSTG | -1.04 | 223 | 0.300 | 0.410 | -0.98 | 223 | 0.329 | 0.479 |
| 24 | RSTG | -1.16 | 195 | 0.249 | 0.355 | -0.79 | 195 | 0.430 | 0.587 |
| 25 | RSTG | -0.89 | 223 | 0.377 | 0.489 | -0.80 | 223 | 0.425 | 0.582 |
| 26 | RSTG | -0.47 | 223 | 0.641 | 0.726 | -0.88 | 223 | 0.380 | 0.535 |
| 27 | RSTG | -0.13 | 195 | 0.899 | 0.924 | -0.94 | 195 | 0.349 | 0.500 |
| 28 | RSTG | 0.01 | 223 | 0.996 | 0.996 | -0.77 | 223 | 0.442 | 0.599 |
| 29 | RSTG | 0.33 | 223 | 0.744 | 0.805 | -0.69 | 223 | 0.492 | 0.629 |
| 30 | RSTG | 0.28 | 111 | 0.780 | 0.835 | -0.51 | 111 | 0.611 | 0.723 |
| 0 | RMTG | -2.60 | 195 | 0.010 | 0.030* | 4.84 | 195 | 0.000 | < 0.001*** |
| 1 | RMTG | -2.49 | 223 | 0.014 | 0.038* | 4.31 | 223 | 0.000 | < 0.001*** |
| 2 | RMTG | -2.15 | 223 | 0.033 | 0.080 | 3.42 | 223 | 0.001 | 0.004** |
| 3 | RMTG | -1.93 | 195 | 0.056 | 0.115 | 2.77 | 195 | 0.006 | 0.023* |
| 4 | RMTG | -2.23 | 223 | 0.027 | 0.067 | 2.91 | 223 | 0.004 | 0.016* |
| 5 | RMTG | -2.73 | 195 | 0.007 | 0.023* | 2.38 | 195 | 0.018 | 0.054 |
| 6 | RMTG | -3.93 | 223 | 0.000 | < 0.001*** | 2.19 | 223 | 0.030 | 0.077 |
| 7 | RMTG | -5.00 | 223 | 0.000 | < 0.001*** | 1.67 | 223 | 0.096 | 0.190 |
| 8 | RMTG | -5.65 | 195 | 0.000 | < 0.001*** | 1.09 | 195 | 0.279 | 0.424 |
| 9 | RMTG | -7.19 | 223 | 0.000 | < 0.001*** | 0.73 | 223 | 0.468 | 0.612 |
| 10 | RMTG | -7.79 | 223 | 0.000 | < 0.001*** | 0.81 | 223 | 0.420 | 0.576 |
| 11 | RMTG | -7.11 | 195 | 0.000 | < 0.001*** | 1.17 | 195 | 0.244 | 0.383 |
| 12 | RMTG | -6.98 | 223 | 0.000 | < 0.001*** | 1.88 | 223 | 0.061 | 0.133 |
| 13 | RMTG | -5.23 | 195 | 0.000 | < 0.001*** | 2.39 | 195 | 0.018 | 0.054 |
| 14 | RMTG | -3.52 | 223 | 0.001 | 0.002** | 3.29 | 223 | 0.001 | 0.006** |
| 15 | RMTG | -1.38 | 223 | 0.168 | 0.267 | 3.76 | 223 | 0.000 | 0.002** |
| 16 | RMTG | 0.15 | 195 | 0.879 | 0.908 | 3.69 | 195 | 0.000 | 0.002** |
| 17 | RMTG | 1.02 | 223 | 0.311 | 0.419 | 3.59 | 223 | 0.000 | 0.003** |
| 18 | RMTG | 1.05 | 223 | 0.293 | 0.403 | 2.96 | 223 | 0.003 | 0.014* |
| 19 | RMTG | 0.61 | 195 | 0.542 | 0.649 | 2.24 | 195 | 0.026 | 0.070 |
| 20 | RMTG | 0.04 | 223 | 0.965 | 0.976 | 1.98 | 223 | 0.049 | 0.112 |
| 21 | RMTG | -0.65 | 223 | 0.513 | 0.624 | 1.50 | 223 | 0.135 | 0.243 |
| 22 | RMTG | -0.90 | 195 | 0.372 | 0.484 | 1.14 | 195 | 0.257 | 0.398 |
| 23 | RMTG | -1.04 | 223 | 0.301 | 0.410 | 1.12 | 223 | 0.265 | 0.406 |
| 24 | RMTG | -1.32 | 195 | 0.189 | 0.290 | 1.34 | 195 | 0.180 | 0.298 |
| 25 | RMTG | -1.63 | 223 | 0.104 | 0.185 | 1.57 | 223 | 0.118 | 0.221 |
| 26 | RMTG | -1.76 | 223 | 0.081 | 0.153 | 1.41 | 223 | 0.161 | 0.276 |
| 27 | RMTG | -1.46 | 195 | 0.147 | 0.241 | 0.75 | 195 | 0.453 | 0.606 |
| 28 | RMTG | -1.45 | 223 | 0.148 | 0.242 | 0.65 | 223 | 0.517 | 0.644 |
| 29 | RMTG | -0.16 | 223 | 0.871 | 0.904 | -0.30 | 223 | 0.765 | 0.849 |
| 30 | RMTG | 0.24 | 111 | 0.813 | 0.859 | -0.59 | 111 | 0.560 | 0.680 |
| 0 | RSMG | -0.78 | 195 | 0.439 | 0.553 | 1.43 | 195 | 0.154 | 0.267 |
| 1 | RSMG | -1.20 | 223 | 0.232 | 0.335 | 1.94 | 223 | 0.053 | 0.117 |
| 2 | RSMG | -1.75 | 223 | 0.082 | 0.154 | 1.95 | 223 | 0.052 | 0.116 |
| 3 | RSMG | -2.22 | 195 | 0.028 | 0.069 | 1.96 | 195 | 0.051 | 0.115 |
| 4 | RSMG | -2.95 | 223 | 0.004 | 0.012* | 1.95 | 223 | 0.053 | 0.117 |
| 5 | RSMG | -3.20 | 195 | 0.002 | 0.006** | 1.26 | 195 | 0.208 | 0.337 |
| 6 | RSMG | -3.72 | 223 | 0.000 | 0.001** | 0.69 | 223 | 0.492 | 0.629 |
| 7 | RSMG | -3.79 | 223 | 0.000 | < 0.001*** | -0.34 | 223 | 0.733 | 0.825 |
| 8 | RSMG | -3.66 | 195 | 0.000 | 0.001** | -1.62 | 195 | 0.107 | 0.206 |
| 9 | RSMG | -4.24 | 223 | 0.000 | < 0.001*** | -3.07 | 223 | 0.002 | 0.011* |
| 10 | RSMG | -4.47 | 223 | 0.000 | < 0.001*** | -3.52 | 223 | 0.001 | 0.003** |
| 11 | RSMG | -4.23 | 195 | 0.000 | < 0.001*** | -3.39 | 195 | 0.001 | 0.005** |
| 12 | RSMG | -4.64 | 223 | 0.000 | < 0.001*** | -3.83 | 223 | 0.000 | 0.001** |
| 13 | RSMG | -4.12 | 195 | 0.000 | < 0.001*** | -3.80 | 195 | 0.000 | 0.001** |
| 14 | RSMG | -3.71 | 223 | 0.000 | 0.001** | -3.97 | 223 | 0.000 | < 0.001*** |
| 15 | RSMG | -2.38 | 223 | 0.018 | 0.049* | -3.62 | 223 | 0.000 | 0.002** |
| 16 | RSMG | -0.60 | 195 | 0.546 | 0.652 | -2.97 | 195 | 0.003 | 0.014* |
| 17 | RSMG | 1.06 | 223 | 0.291 | 0.401 | -2.98 | 223 | 0.003 | 0.014* |
| 18 | RSMG | 2.13 | 223 | 0.035 | 0.082 | -3.03 | 223 | 0.003 | 0.012* |
| 19 | RSMG | 2.27 | 195 | 0.024 | 0.061 | -3.00 | 195 | 0.003 | 0.013* |
| 20 | RSMG | 2.42 | 223 | 0.016 | 0.045* | -3.30 | 223 | 0.001 | 0.006** |
| 21 | RSMG | 2.00 | 223 | 0.047 | 0.102 | -3.50 | 223 | 0.001 | 0.003** |
| 22 | RSMG | 1.64 | 195 | 0.102 | 0.182 | -3.19 | 195 | 0.002 | 0.008** |
| 23 | RSMG | 1.68 | 223 | 0.095 | 0.175 | -2.86 | 223 | 0.005 | 0.018* |
| 24 | RSMG | 1.58 | 195 | 0.117 | 0.202 | -1.44 | 195 | 0.153 | 0.267 |
| 25 | RSMG | 1.62 | 223 | 0.107 | 0.189 | 0.17 | 223 | 0.861 | 0.921 |
| 26 | RSMG | 1.37 | 223 | 0.172 | 0.272 | 1.45 | 223 | 0.149 | 0.262 |
| 27 | RSMG | 0.65 | 195 | 0.515 | 0.624 | 1.48 | 195 | 0.140 | 0.249 |
| 28 | RSMG | -0.26 | 223 | 0.797 | 0.847 | 1.67 | 223 | 0.096 | 0.190 |
| 29 | RSMG | -1.00 | 223 | 0.317 | 0.426 | 0.76 | 223 | 0.446 | 0.601 |
| 30 | RSMG | -1.15 | 111 | 0.254 | 0.361 | 0.12 | 111 | 0.907 | 0.942 |
| 0 | RAG | 2.13 | 188 | 0.034 | 0.082 | 0.57 | 188 | 0.567 | 0.682 |
| 1 | RAG | 1.25 | 215 | 0.211 | 0.310 | -0.15 | 215 | 0.877 | 0.924 |
| 2 | RAG | 1.95 | 215 | 0.053 | 0.111 | -0.66 | 215 | 0.512 | 0.644 |
| 3 | RAG | 1.83 | 188 | 0.069 | 0.137 | -0.67 | 188 | 0.507 | 0.639 |
| 4 | RAG | 2.07 | 215 | 0.040 | 0.091 | -0.63 | 215 | 0.530 | 0.657 |
| 5 | RAG | 1.79 | 188 | 0.076 | 0.145 | -1.06 | 188 | 0.290 | 0.434 |
| 6 | RAG | 1.63 | 215 | 0.105 | 0.186 | -1.76 | 215 | 0.079 | 0.165 |
| 7 | RAG | 1.41 | 215 | 0.161 | 0.258 | -2.57 | 215 | 0.011 | 0.036* |
| 8 | RAG | 1.64 | 188 | 0.102 | 0.182 | -2.61 | 188 | 0.010 | 0.034* |
| 9 | RAG | 2.06 | 215 | 0.041 | 0.092 | -2.56 | 215 | 0.011 | 0.037* |
| 10 | RAG | 2.61 | 215 | 0.010 | 0.029* | -1.65 | 215 | 0.100 | 0.195 |
| 11 | RAG | 3.14 | 188 | 0.002 | 0.008** | -0.68 | 188 | 0.495 | 0.630 |
| 12 | RAG | 4.31 | 215 | 0.000 | < 0.001*** | 0.16 | 215 | 0.869 | 0.924 |
| 13 | RAG | 5.15 | 188 | 0.000 | < 0.001*** | 0.85 | 188 | 0.396 | 0.551 |
| 14 | RAG | 6.71 | 215 | 0.000 | < 0.001*** | 1.70 | 215 | 0.090 | 0.182 |
| 15 | RAG | 7.70 | 215 | 0.000 | < 0.001*** | 2.39 | 215 | 0.018 | 0.053 |
| 16 | RAG | 8.01 | 188 | 0.000 | < 0.001*** | 2.70 | 188 | 0.007 | 0.027* |
| 17 | RAG | 9.39 | 215 | 0.000 | < 0.001*** | 2.74 | 215 | 0.007 | 0.025* |
| 18 | RAG | 9.76 | 215 | 0.000 | < 0.001*** | 1.87 | 215 | 0.062 | 0.135 |
| 19 | RAG | 9.11 | 188 | 0.000 | < 0.001*** | 0.74 | 188 | 0.459 | 0.607 |
| 20 | RAG | 9.44 | 215 | 0.000 | < 0.001*** | 0.08 | 215 | 0.934 | 0.959 |
| 21 | RAG | 9.08 | 215 | 0.000 | < 0.001*** | -0.46 | 215 | 0.643 | 0.750 |
| 22 | RAG | 8.08 | 188 | 0.000 | < 0.001*** | -0.57 | 188 | 0.570 | 0.684 |
| 23 | RAG | 8.00 | 215 | 0.000 | < 0.001*** | -0.59 | 215 | 0.553 | 0.674 |
| 24 | RAG | 6.91 | 188 | 0.000 | < 0.001*** | -0.02 | 188 | 0.983 | 0.994 |
| 25 | RAG | 6.81 | 215 | 0.000 | < 0.001*** | 0.51 | 215 | 0.608 | 0.722 |
| 26 | RAG | 6.13 | 215 | 0.000 | < 0.001*** | 0.88 | 215 | 0.380 | 0.535 |
| 27 | RAG | 4.82 | 188 | 0.000 | < 0.001*** | 0.34 | 188 | 0.737 | 0.828 |
| 28 | RAG | 4.51 | 215 | 0.000 | < 0.001*** | -0.06 | 215 | 0.949 | 0.967 |
| 29 | RAG | 4.30 | 215 | 0.000 | < 0.001*** | -1.59 | 215 | 0.112 | 0.215 |
| 30 | RAG | 3.10 | 107 | 0.002 | 0.009** | -1.57 | 107 | 0.119 | 0.221 |

Notes: Positive *t*-statistics in the MET > LIT comparisons indicate tests where activation was greater for metaphorical compared to literal sentences whereas negative *t*-statistics indicate the opposite. Similarly positive *t*-statistics in the NOV/MET > FAM/MET comparisons indicate tests where activation was greater for novel compared to familiar metaphors whereas negative *t*-statistics indicate the opposite direction of the effects. Correction for multiple comparisons was conducted across regions of interest (ROIs) and 1s windows for each contrast at a false discovery rate (FDR) where **q* < 0.05, ***q* < 0.01, and ****q* < 0.001. Abbreviations: L = left; R = right; SFG = superior frontal gyrus; MFG = middle frontal gyrus; IFGtri = inferior frontal gyrus, pars triangularis; IFGop = IFG, pars opercularis; STG = superior temporal gyrus; MTG = middle temporal gyrus; ITG = inferior temporal gyrus; SMG = supramarginal gyrus; AG = angular gyrus.
